# GLP-1R agonists activate human hypothalamic neurons

**DOI:** 10.1101/2024.04.02.587825

**Authors:** Simone Mazzaferro, Hsiao-Jou Cortina Chen, Olivier Cahn, Andrian Yang, Dmytro Shepilov, Eugene Seah, Jiahui Chen, Bavesh Jawahar, Constanza Alcaino, Viviana Macarelli, Iman D. Mali, John A. Tadross, Fiona Gribble, Frank Reimann, John C. Marioni, Florian T. Merkle

## Abstract

Drugs like semaglutide (a.k.a. Ozempic/Wegovy) that activate the glucagon-like peptide-1 receptor (GLP-1R) are a promising therapy for obesity and type 2 diabetes (T2D). Animal studies suggest that these drugs likely function by stimulating GLP-1R on appetite-suppressing neuron populations in the brain, but it is still unclear how they act to reduce food intake in humans. We therefore generated appetite-regulatory hypothalamic neurons from human pluripotent stem cells (hPSCs) to study their responses to GLP-1R agonists by calcium imaging and electrophysiology. We found that hPSC-derived proopiomelanocortin (POMC) and other hypothalamic neuron subtypes expressed *GLP1R* mRNA, and many of these neurons robustly responded to GLP-1R agonists by membrane depolarization, increased action potential firing, and extracellular calcium influx that persisted long after agonist withdrawal. The observed GLP-1R-induced response was likely mediated by the activation of PKA and L-type calcium channels, and led to significant changes in gene expression. These findings provide mechanistic insight into how GLP-1R agonists may suppress appetite in humans.

## Introduction

Obesity is a condition of excess adiposity that affects over a billion people worldwide and significantly increases the risk of chronic diseases such as type 2 diabetes (T2D) and cardiovascular disease (NCD Risk Factor Collaboration (NCD-RisC) 2024). Obesity has a strong genetic basis, and genetic variants associated with obesity predominantly act in the brain (Locke et al. 2015). Studies of individuals with severe, early-onset obesity revealed mutations in brain circuits that are essential for appetite regulation, such as the leptin-melanocortin system (van der Klaauw et al. 2019; Farooqi and O’Rahilly 2006; Loos and Yeo 2022). Specifically, populations of pro-opiomelanocortin (POMC) neurons in the arcuate nucleus of the hypothalamus increase their firing rate in response to the adipocyte-derived hormone leptin (Cowley et al. 2001; Qiu et al. 2018) and the predominantly gut-derived hormone glucagon-like peptide-1 (GLP-1) (Gabery et al. 2020; Secher et al. 2014; Dong et al. 2021; Péterfi et al. 2021). Activated POMC neurons then release the POMC-derived peptides α-melanocyte stimulating hormone (α-MSH) and β-MSH (Kirwan et al. 2018; Lee et al. 2006) that stimulate the melanocortin 4 receptor (MC4R) on neurons that suppress food intake and increase energy expenditure (Fenselau et al. 2017; Baldini and Phelan 2019; Cone 2005). The loss of either *POMC* or *MC4R* is sufficient to increase appetite and body weight in both humans (Farooqi et al. 2003; Yeo et al. 2003; Krude et al. 1998) and animals (Dittmann et al. 2024; Yaswen et al. 1999; Huszar et al. 1997). Other key brain cell populations that respond to GLP-1 include neurons in the area postrema (AP) (Kawatani et al. 2018) and nucleus tractus solitarius (NTS) of the hindbrain (Hayes et al. 2011) that are important for its appetite-suppressing effects in rodents (Alhadeff et al. 2012; Huang et al. 2024).

The ability of GLP-1 to suppress appetite has recently been harnessed by synthetic agonists of the GLP-1 receptor (GLP-1R) that are revolutionizing the treatment of T2D and obesity (Meier 2012; Bessesen and Van Gaal 2018; Tan et al. 2022). GLP-1R agonists delay gastric emptying and reduce food intake in humans to reduce body weight and improve metabolic health for long periods of time (Halawi et al. 2017). The efficacy of GLP-1R agonist monotherapies such as semaglutide can be further improved by combining it with agonism of the gastric inhibitory polypeptide receptor (GIPR) and/or the glucagon receptor (GCGR) (Knerr et al. 2022; Coskun et al. 2022; Zhao et al. 2022), but their efficacy does not yet match the weight loss seen with bariatric surgery and some users experience side effects such as nausea.

To rationally design more effective treatments for obesity, it is important to understand the mechanisms by which existing treatments work in humans. GLP-1R signaling has been well-studied in pancreatic β-cells (Shilleh et al. 2024), where its coupling to Gα_s_ stimulates adenylate cyclase to elevate the intracellular concentration of cyclic adenosine monophosphate, or [cAMP]_i_. This cAMP increase in turn activates protein kinase A (PKA) which phosphorylates residues on L-type voltage-gated calcium channels (VGCCs) to increase the influx of extracellular Ca^2+^ and promote insulin secretion (Gomez et al. 2002). GLP-1R can also couple to β-arrestin upon ligand binding, which promotes its rapid internalization (Sonoda et al. 2008) and continued signaling from the endosome (Irannejad et al. 2013). In the nervous system, signaling pathways downstream of GLP-1R are less well understood but likely also involve Gα_s_ coupling, since PKA becomes activated in NTS neurons upon GLP-1R agonist administration (Hayes et al. 2011). Due to the inaccessibility of human neurons, endogenous neuronal GLP-1R signaling pathways are unknown, and extrapolating insights from rodent studies is complicated by species-specific functional differences in appetite-regulatory cell types (Steuernagel et al. 2022; Kirwan et al. 2018; Tadross et al. 2025).

To address these limitations, we derived hypothalamic neurons from human pluripotent stem cells (hPSCs), including *POMC-GFP* and *POMC-NeonGreen* knock-in reporter cell lines generated on two distinct genetic backgrounds to enable the prospective identification of POMC neurons in live cultures. We found that many human hypothalamic neurons expressed *GLP1R* mRNA and were significantly activated by GLP-1 and other GLP-1R agonists including exendin-4 derivatives, liraglutide, semaglutide, and the dual GLP-1R/GIPR agonist tirzepatide (a.k.a. Mounjaro). This activation was completely blocked by the specific GLP-1R antagonist exendin-(9-39). Electrophysiological recordings and pharmacological studies showed that POMC neurons were depolarized by GLP-1R agonists for at least 20 minutes and fired substantially more action potentials via mechanisms that required functional L-type calcium channels. Prolonged stimulation with semaglutide also induced significant transcriptional changes in human POMC neurons related to intracellular calcium signaling, cell survival, and neurodegeneration. Together, these studies demonstrate that appetite-regulatory human neurons are durably activated by GLP-1R agonists via similar mechanisms as those reported in pancreatic β-cells, and suggest that the activation of POMC or other hypothalamic neurons may contribute to the observed appetite suppressive effects of GLP-1R agonists drugs in humans.

## Results

### Human hypothalamic neurons express *GLP1R*

To investigate GLP-1R signaling in human hypothalamic and POMC neurons, we differentiated two genetically distinct hPSC lines (HUES9 and KOLF2.1J) into hypothalamic cells using established protocols (Chen, Mazzaferro, et al. 2023; Merkle et al. 2015; Kirwan et al. 2017). Differentiated cultures contained large numbers of neurons characterized by extensive neurite outgrowth (Fig. 1A, S1E), and a subpopulation of cultures were strongly immunoreactive for POMC/α-MSH (Fig. 1B, S1F). To assess whether hPSC-derived hypothalamic neurons express *GLP1R* mRNA, we performed RNAscope multiplex fluorescent in situ hybridization for *GLP1R* and *POMC* transcripts. We found that both *POMC*+ and *POMC*-cells contained puncta corresponding to *GLP1R* mRNA (Fig. 1C, Fig. S1G). Quantitative analysis of these images revealed that within HUES9-derived hypothalamic neurons, 32% (353/754) of *POMC*+ neurons expressed *GLP1R* whereas only 14% (5880/37286) of POMC-neurons were *GLP1R*+ (Fig. 1D). Similarly, within KOLF2.1J-derived hypothalamic neurons, 23% (93/320) of *POMC*+ neurons expressed *GLP1R* whereas only 8% (496/5624) of *POMC*-neurons were *GLP1R*+ (Fig. S1H). Furthermore, within the *GLP1R*+ populations the number of *GLP1R* puncta per neuron was significantly higher in the *POMC*+ than in *POMC*-neurons in both cell lines (p<0.05) suggesting higher overall receptor expression levels in *POMC*+ cells (Fig. 1E; S1I; Table S1A).

**Figure 1.**
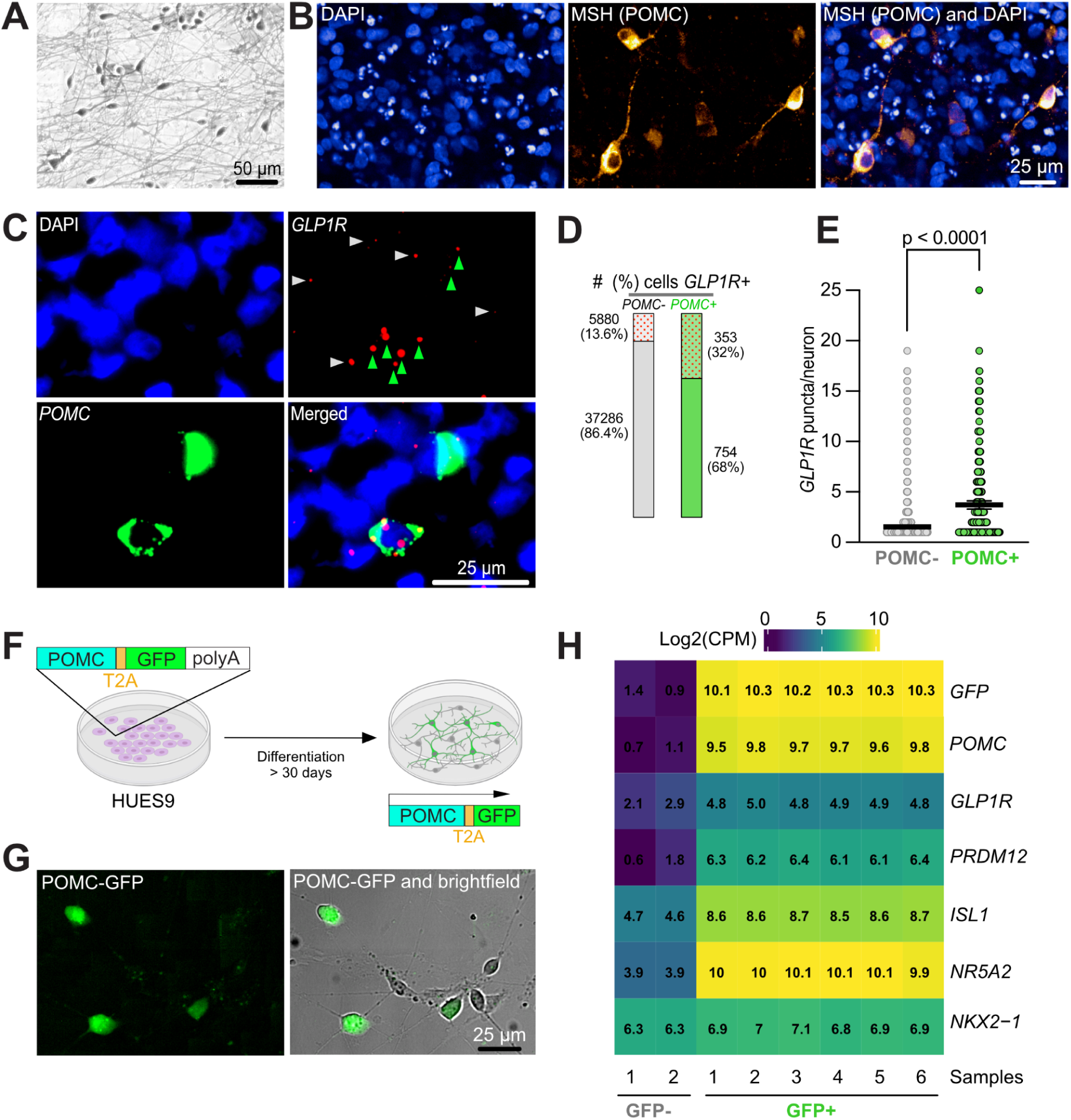
Expression of *GLP1R* mRNA in human POMC neurons. A,B) Human pluripotent stem cells were differentiated into hypothalamic neurons that exhibited mature neuronal morphology with extensive neurite outgrowth (A) and contained POMC neurons, as confirmed by immunostaining for α-MSH (B). **C**) Photomicrographs of RNAscope fluorescent in situ hybridization showing *GLP1R* mRNA puncta in both POMC+ and POMC-cells. **D,E**) *GLP1R* puncta were detected in a larger fraction of POMC+ neurons (353/1107 cells, 32%) than POMC-neurons (5880/43166 cells, 13.6%, D), and POMC+ neurons also contained, on average, a higher number of *GLP1R* puncta per cell (POMC+: 3.7 puncta/cell, n=353; POMC-: 1.5 puncta/cell, n=5880; p<0.001, unpaired Mann-Whitney test, E). **F,G**) To confirm validate POMC neuron identity, a POMC-T2A-GFP reporter hPSC line was used to generate hypothalamic neurons (F), in which GFP+ cells could be readily identified in live cultures (G). **H**) Transcriptomic analysis of FACS-purified GFP-and GFP+ populations confirmed enrichment of *GLP1R* and genes associated with POMC neuron identity in GFP+ cells. Scale bars, 25 µm.

**Figure S1.**
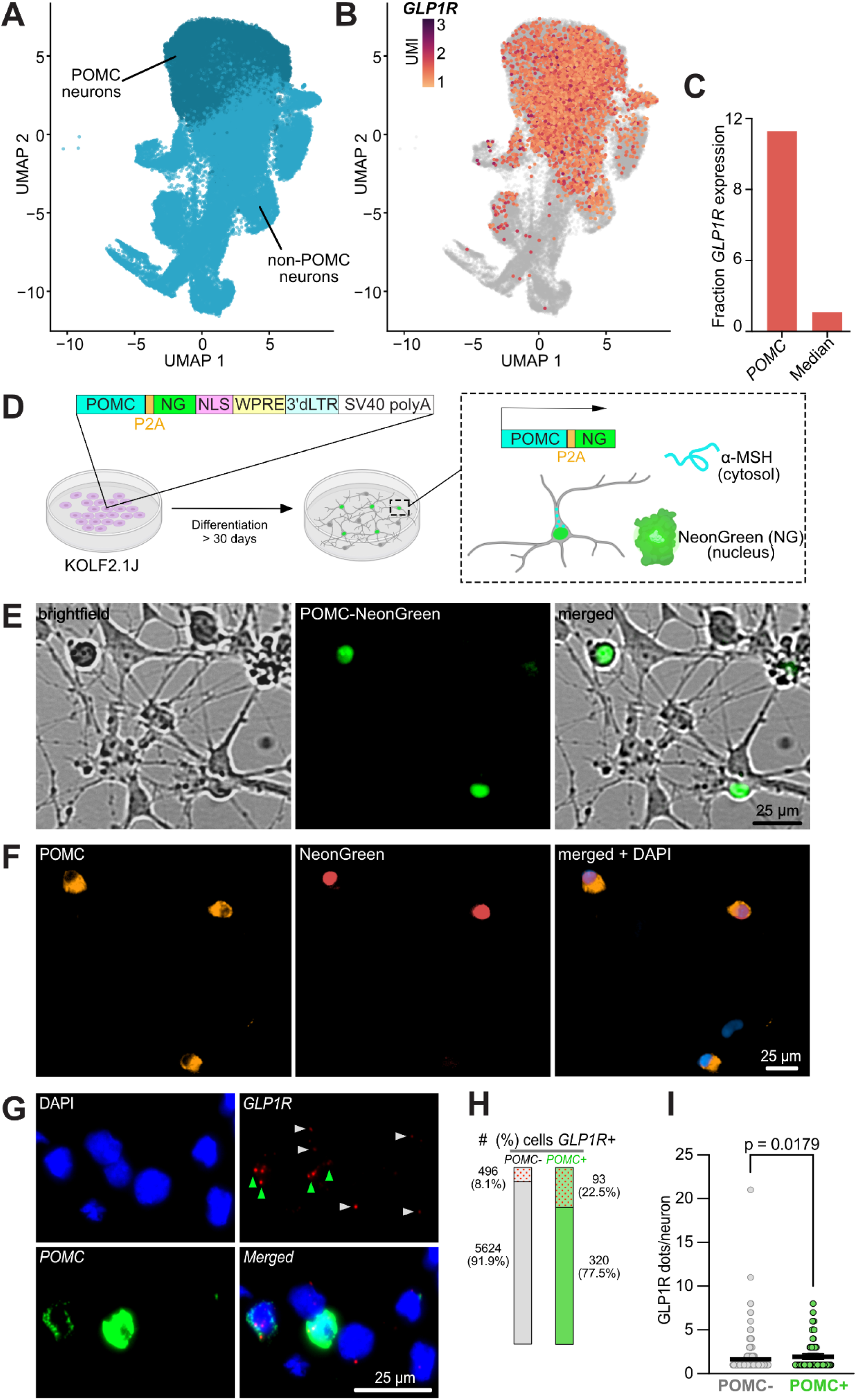
*GLP1R* expression in a KOLF2.1J-POMC-NG reporter cell line. **A-C**) UMAP of single-cell RNA sequencing data from hPSC-derived hypothalamic POMC (dark blue) or non-POMC (light blue) neurons (A) showing expression of *GLP1R* mRNA (UMI≥1) across hypothalamic neuron populations (B) and relative enrichment within POMC neuron clusters (2578/22892 cells, 11.3%) compared with a median of 1.1% across all other clusters (C). **D**) To validate POMC neuron identity in an independent genetic background, a KOLF2.1J iPSC line was engineered by replacing the endogenous *POMC* stop codon with an in-frame construct containing a self-cleaving P2A peptide followed by NeonGreen (NG) fused to a nuclear localization signal (NLS) and the post-transcriptional regulatory elements WPRE, 3′dLTR, and SV40 polyA to drive nuclear green fluorescence in POMC neurons. **E,F**) Upon hypothalamic differentiation, nuclear NeonGreen fluorescence was readily detectable in live POMC neurons (E) that overlapped with POMC immunoreactivity upon immunostaining (F). **G-I**) RNAscope fluorescent in situ hybridization for *POMC* and *GLP1R* in this reporter line revealed *GLP1R* expression in both POMC+ and POMC-cells (G) with *GLP1R* present in larger fraction of POMC+ cells (93/413 cells, 22.5%) than POMC-cells (496/6120 cells, 8.1%) (H), and POMC+ neurons contained more *GLP1R* puncta per cell (POMC+: 1.9 puncta/cell, n=93; POMC-: 1.6 puncta/cell, n=496; p=0.018, unpaired Mann-Whitney test) (I). Scale bars, 25 μm.

To test the generalizability of these findings, we analyzed single-cell RNA sequencing data from hypothalamic neurons differentiated from multiple hPSC lines (Chen, Yang, et al. 2023) and found that *GLP1R* transcript was detectable (UMI ≥ 1) in a substantial fraction of cells in clusters annotated as POMC neurons (2578/22892 cells, 11.3%), whereas a median of 1.1% of neurons in other clusters expressed *GLP1R* (Fig. S1A-C; Table S1E). These findings closely resemble those seen in an integrated single-cell and single-nucleus RNAseq atlas of mouse hypothalamic cells (Steuernagel et al. 2022), where *Glp1r* was expressed in 380/4655 (8.2%) of cells in a cluster (C66-19) annotated as *Pomc* neurons, while the median percentage of *Glp1r*-expressing cells in other clusters was 1.1%. These percentages are likely underestimates due to the imperfect transcript detection efficiency of RNAscope and scRNAseq (Brennecke et al. 2013), and since functional studies have found that 37-100% of rodent Pomc neurons respond to GLP-1R agonists (Rønnekleiv et al. 2014; He et al. 2019; Péterfi et al. 2021; Secher et al. 2014).

### Generation of POMC reporter cell lines

To study GLP-1R signaling in human POMC and other hypothalamic neurons, we generated a POMC-GFP knock-in cell line on the HUES9 genetic background (Fig. 1F) (Chen, Yang, et al. 2023) as well as a POMC-NeonGreen (NG) knock-in cell line (Fig. S1D) on the KOLF2.1J genetic background (Pantazis et al. 2022). In both cases, we designed these reporters to preserve functional endogenous POMC expression followed by 2A peptides to induce ribosomal skipping and co-translationally express the fluorescent reporter gene (Fig. 1G; S1E). The NeonGreen reporter gene contained a nuclear localization sequence (NLS) to concentrate fluorescence to the nucleus. To confirm the fidelity of these reporter lines, we flow-purified fluorescent cells and immunostained for POMC (Chen, Yang, et al. 2023), and also characterized GFP+ and GFP-cells by bulk RNA sequencing (Fig. 1H). This transcriptomic analysis showed that GFP+ neurons expressed not just POMC and the reporter gene, but also *GLP1R* and transcription factors involved in POMC neuron differentiation including *PRDM12*, *ISL1*, *NR5A2*, and *NKX2-1* (Hael et al. 2020; Yu et al. 2022; Orquera et al. 2019) (Fig. 1H, Table S1D). Live imaging confirmed that POMC neurons could be prospectively identified and studied by fluorescent microscopy in both the POMC-GFP (Fig. 1G) and POMC-NG cell lines (Fig. S1F). Together, these results demonstrate that human stem-cell derived hypothalamic neurons express *GLP1R* transcript across multiple reporter lines, establishing a scalable cellular model for investigating GLP-1R signaling in appetite-regulatory human neurons.

### GLP-1R agonists increase intracellular calcium in human hypothalamic neurons

The expression of *GLP1R* on human POMC and other hypothalamic neurons suggested that they might functionally respond to GLP-1 peptide and other GLP-1R agonists. Since these agonists increase the electrical activity of rodent POMC neurons (Péterfi et al. 2021; Secher et al. 2014; Dong et al. 2021; Rønnekleiv et al. 2014; He et al. 2019; Singh et al. 2022) and raise their intracellular calcium concentration ([Ca^2+^]_i_), we performed calcium imaging on human hypothalamic neurons derived from the *POMC-GFP* reporter line by loading them with a red calcium-sensitive dye (Fig. 2A). To isolate cell-autonomous responses from network activity, we included synaptic blockers in all media and solutions during recording sessions..

**Figure 2.**
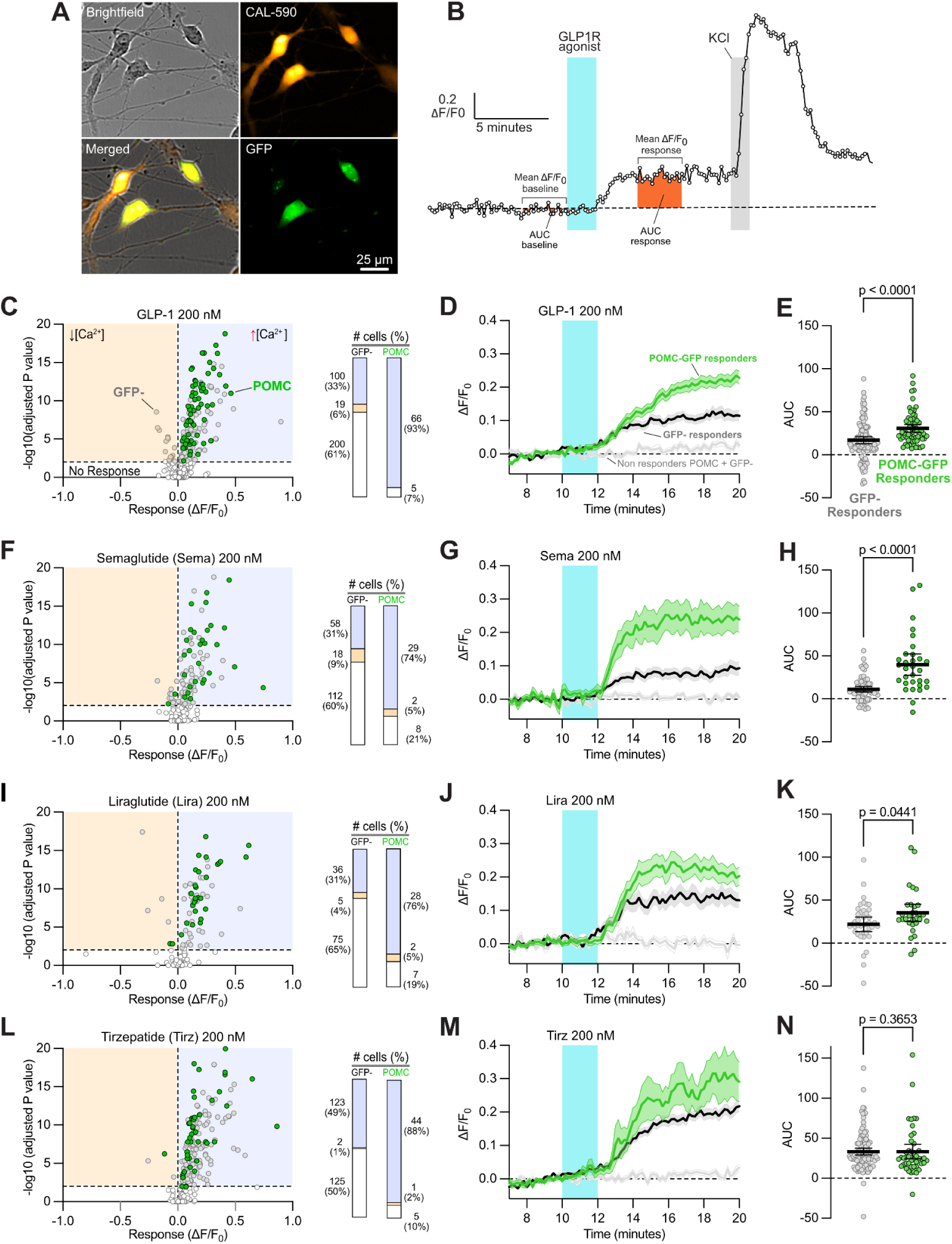
A subset of human hypothalamic neurons respond to GLP-1R agonists. **A)** Photomicrographs from a *POMC-GFP* reporter hPSC line showing neuronal morphology in brightfield, baseline fluorescence from the Cal-590 AM calcium-sensitive dye, endogenous GFP fluorescence from the reporter, and a merge of these images. **B)** Experimental schematic for calcium imaging, where normalized changes in calcium indicator fluorescence (Y axis, ΔF/F_0_) over time (X axis, minutes) are measured before and after administration of a candidate factor (e.g. GLP-1R agonist, blue bar) followed by cell depolarization with 50 mM potassium chloride (KCl, grey bar) to identify cells that significantly respond to the candidate factor based on the area under the curve (AUC) and/or difference in mean response amplitude over baseline. **C)** Graph of response magnitude (X axis, ΔF/F_0_) vs statistically significance of response (Y axis, p value from paired t-tests) to 200 nM recombinant human GLP-1 showing the classification of GFP-cells ((n=319, grey) or GFP+ POMC cells (n=71, green) into categories of significantly inhibited (light orange) significantly activated (light blue) or no significant response (white). Among responding cells (bar graphs, right) the proportion of POMC neurons activated by GLP-1 (66/71, 93%) was significantly higher (p<0.0001, Chi-square test) than GFP-cells (100/319, 33%). **D)** Average kinetics after administration of GLP-1 among responding GFP+ cells (green), responding GFP-cells (black and grey) and non-responding cells (light grey) with error bars indicating SEM (lighter shading). **E)** Distribution of response magnitudes (AUC) among all responder cells was significantly higher (p<0.0001; unpaired Mann-Whitney t-test) in POMC responders than GFP-responders. **F-N)** Similar analysis as shown in C-E was performed with 200 nM GLP-1R agonists semaglutide (F), liraglutide (I) or the GLP-1R/GIPR dual agonist drug tirzepatide (L), showing that the proportion of POMC neurons activated by these ligands was significantly higher than for GFP-cells (p<0.0001, Chi-square test). Response magnitudes among responder cells were significantly higher (unpaired Mann-Whitney t-test) than GFP-cells for semaglutide (H; p<0.0001) and liraglutide (K; p<0.05) but not for tirzepatide (N; p=0.37).

**Figure S2.**
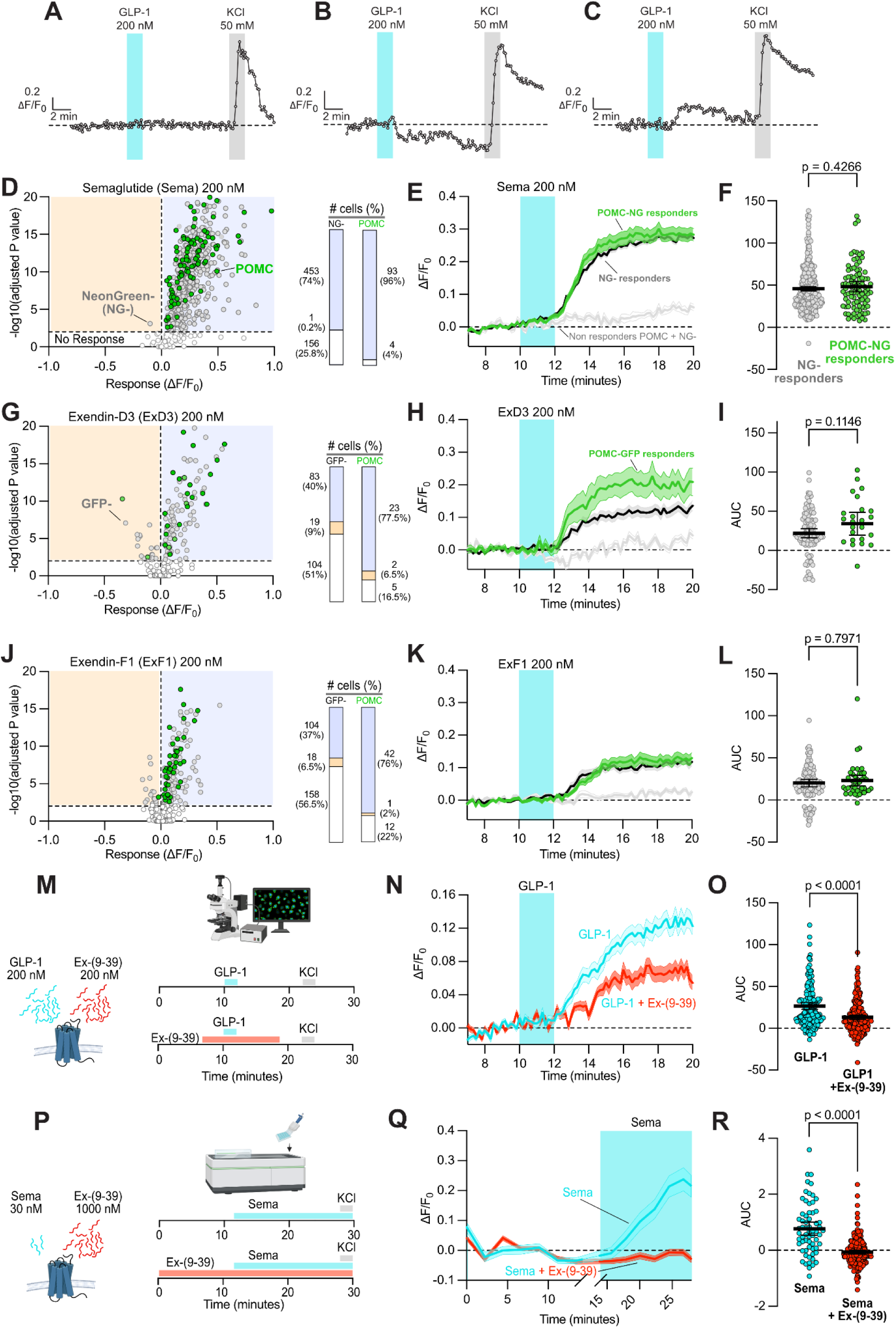
GLP-1R agonist responses are consistent in different reporter lines and are blocked by GLP-1R antagonists. **A-C)** Representative calcium imaging traces showing cells that lack detectable responses (A), show apparent inhibitory responses despite the presence of synaptic blockers (B) or show excitatory responses (C) to 200 nM GLP-1. **D)** Response amplitude vs response significance of NG+ POMC neurons (n=96) and NG-neurons (n=610) to 200 nM semaglutide, revealing that among responders (D, bar graphs) the proportion of POMC neurons activated (75/96, 78%) by 200 nM semaglutide was significantly higher (p<0.0001, Chi-square test) than for NG-neurons (100/319, 33%). Axes are as in Fig. 2C. **E,F**) The average traces of responding cells show that the calcium indicator fluorescence increases and persists after semaglutide administration (E) to a significantly greater (p<0.05) extent in NG+ cells than NG-cells (F). **G-L)** Analysis follows the same format as shown in (D-F), with significantly more (p<0.0001, Chi-square test) GFP+ POMC neurons (25/30, 83%) than GFP-neurons (102/206, 49%) responding to both exendin-D3 (G) and extendin-F1 (J; POMC neurons: 43/55, 78%; GFP-cells 122/280, 44%), but showing no significant differences in response magnitude to extendin-D3 (H,I; n= 25 POMC neurons and n=102 GFP-cells), or to exendin-F1 (K,L; n=42 POMC neurons and n=122 GFP-cells) where response magnitudes tended to be lower. **M-O**) Co-administration of equimolar ratios of GLP-1 and the GLP-1R antagonist exendin-(9-39) (Ex-(9-39)) as schematized in (M) significantly reduced (p<0.0001, unpaired Mann-Whitney t-test) neuronal responses by approximately 50% (N) as quantified by AUC calculations (O). N=178 GLP-1 treated, and n=206 GLP-1 and exendin-(9-39)-treated neurons. **P-R)** Excess concentrations exendin-(9-39) as schematized in (P) were sufficient to completely block (p<0.0001, unpaired Mann-Whitney t-test) POMC-GFP+ neuron responses to 30 nM semaglutide (Q), as quantified by AUC calculations (R). N=64 semaglutide-treated and n=200 semaglutide + exendin-(9-39)-treated GFP+ neurons.

After 10 minutes of recording to identify cells with stable baseline fluorescence, we added 200 nM GLP-1 to cultures for two minutes followed by a wash-out period of 10-20 minutes and a terminal application of 50 mM potassium chloride (KCl) to identify neurons capable of significant depolarization-mediated calcium influx. The concentration of GLP-1R was within the range (100 nM - 1 µM) previously used in recordings from rodent slice cultures (He et al. 2019; Péterfi et al. 2021; Secher et al. 2014; Gabery et al. 2020; Dong et al. 2021). We selected neurons for further analysis if they exhibited stable baseline fluorescence and showed a clear increase in fluorescence (ΔF/F_0_) in response to KCl (Fig. 2B; Table S2) which was true for the vast majority of recorded cells. Among analyzed neurons, a proportion showed no response to GLP-1 (Fig. S2A), rare neurons showed reduced calcium indicator dye fluorescence (Fig. S2B), and many neurons showed increased fluorescence intensity that persisted for tens of minutes (Fig. S2C), suggesting prolonged inhibition or activation, respectively.

To categorize non-responsive, inhibited, and activated neurons in an unbiased manner, we tested for significant differences between fluorescence intensities at 18 time points in a three-minute pre-stimulus (baseline) window just before GLP-1 addition, and a three-minute post-stimulus window after GLP-1 wash-out, at which point responses in activated or inhibited cells had plateaued (Fig. 2B). We defined responsive cells as those having significant (p<0.01, t-test) changes in ΔF/F_0_, and either inhibited or activated depending on the direction of the response, and all other cells as non-responsive (Fig. 2C). Using these criteria, we found that the majority of GFP+ POMC neurons (66/71, 93%) were activated by 200 nM GLP-1. In contrast, fewer (100/319, 33%) GFP-neurons were activated and some GFP-neurons (19/319, 6%) appeared to be inhibited (Fig. 2C). We then quantified the response magnitudes by calculating the post-stimulus area under the curve (AUC) normalized to the pre-stimulus AUC (Fig. 2E) during the same baseline and post-stimulus time windows described above (Fig. 2B; Table S2C-F). We found that the mean response magnitude of responder cells was significantly greater in POMC neurons than in GFP-cells (p<0.0001; Fig. 2D,E; Table SA,B)

Intrigued by the responses of human POMC neurons to GLP-1, we wondered how they would respond to GLP-1R agonist drugs, some of which are clinically used to treat T2D and obesity. We therefore treated hPSC-derived hypothalamic cultures with 200 nM semaglutide (Lau et al. 2015; Wilding et al. 2021), liraglutide (Knudsen and Lau 2019), the biased agonists Exendin-D3 and Exendin-F1 (Fang et al. 2020; Jones et al. 2018; Lucey et al. 2020; Jones 2022) or the dual GLP-1R/GIPR agonist tirzepatide (Coskun et al. 2018; Jastreboff et al. 2022) and compared their responses to those observed with 200 nM GLP-1 (Fig. 2C-E; Table S2A). We found that significantly more (p<0.0001) POMC neurons (31/39, 79%) than GFP-neurons (76/188, 40%) showed a detectable response to these agonists (Fig. 2F; Table S2A). Semaglutide predominantly activated rather than inhibited both POMC neurons (29/31 cells, 94%), and GFP-neurons (58/76 cells, 76%) and produced a significantly greater calcium response (p<0.0001) in POMC neurons than in GFP-(Fig. 2G,H; Table S2A). These findings were corroborated in a genetically distinct POMC-NG reporter line in which the vast majority of both NG+ POMC neurons (93/97, 96%) and NG-neurons (454/610, 74%) showed a detectable and nearly always positive response, with POMC-NG+ more likely to respond than NG-neurons (Fig. S2D; Table S2A; p<0.0001). However, we did not detect a significant difference in the magnitude of semaglutide-induced responses between POMC-NG+ and NG-neurons (Fig. S2E,F; Table S2A; p=0.43).

Liraglutide administration elicited responses in significantly (p<0.0001) more GFP+ POMC neurons (30/37, 84%) than GFP-neurons (41/116, 35%), and the vast majority of both responding POMC neurons (28/30, 93%) and GFP-neurons (36/41, 87%) were activated (Fig. 2I; Table S2A), with POMC neurons showing a greater response magnitude (p<0.05; Fig. 2J,K; Table S2A). We next analyzed the responses of hypothalamic neurons to the ‘biased agonists’ exendin-D3 and exendin-F1 reported to elicit distinct GLP1R signaling and trafficking (Fang et al. 2020; Ben Jones et al. 2018; Lucey et al. 2020) (Fig. S2G-L; Table S2). In response to exendin-D3, significantly (p<0.0001) more POMC neurons (25/30, 83%) than GFP-neurons (102/206, 49%) responded, and among responders the majority of both POMC and GFP-neurons were activated (Fig. S2G; Table S2A). Similarly, significantly (p<0.0001) more POMC neurons (43/55, 78%) than GFP-cells (122/280, 44%) responded to exendin-F1 (Fig. S2J; Table S2A). The magnitude of responses to exendin-D3 (Fig. S2H,I; Table S2) and exendin-F1 (Fig. S2K,L; Table S2A) were not significantly different (p>0.05) between POMC neurons and GFP-neurons. In response to the dual GLP-1R/GIPR agonist tirzepatide, significantly (p<0.0001) more POMC neurons (45/50, 90%) than GFP-neurons (125/250, 50%) responded, and responses were nearly always activating for both POMC neurons (44/45, 98%) and GFP-neurons (123/125, 98%; Fig. 2L; Table S2A) which both responded at similar magnitudes (p>0.05; Fig. 2M,N; Table S2).

To confirm that GLP-1R agonist-induced responses were indeed mediated via GLP-1R activation, we next tested whether the selective GLP-1R antagonist exendin-(9-39) could block GLP-1R agonist-evoked neuronal responses. Indeed, co-application of equimolar concentrations (200 nM) of exendin-(9-39) and GLP-1 (Fig. S2M) significantly reduced (p<0.0001) GLP-1-evoked calcium responses by approximately 50% as expected (Fig. S2N,O, Table S2A), whereas an excess (1 μM) of exendin-(9-39) (Fig. S2P) completely abolished (p<0.0001) responses induced by 30 nM semaglutide (Fig. S2Q,R, Table S2A). These findings demonstrate that engagement of GLP-1R is required for neuronal activation and calcium signaling in response to GLP-1R agonists. Overall, we found that the majority of analyzed POMC neurons and many other hypothalamic neurons responded robustly to a wide variety of GLP-1R agonists, and these responses nearly always increased [Ca^2+^]_i_ and persisted for the duration of the recording (10 minutes) until KCl was added (Fig. 2; Fig. S2, Table S2).

### Semaglutide stimulates human POMC neuron activity

Persistent increases in [Ca^2+^]_i_ could be caused by the Ca^2+^ release from internal stores such as the endoplasmic reticulum, or from neuronal depolarization and the entry of extracellular Ca^2+^ through calcium channels and/or non-selective cation channels. Electrophysiological recordings from hypothalamic mouse brain slices demonstrated that GLP-1 or liraglutide stimulation depolarized POMC neurons in a GLP-1R-dependent manner and that depolarization persisted for at least tens of minutes after agonist removal (Secher et al. 2014; Dong et al. 2021). We therefore hypothesized that the prolonged nature of the calcium signal we observed in calcium imaging experiments of POMC neurons (Fig. 2) corresponded to their depolarization and increased electrophysiological activity.

To test this hypothesis, we performed perforated patch-clamp recordings on POMC neurons derived from both POMG-GFP and POMG-NG reporter lines. We carried out recordings in current-clamp mode at I=0 (no current injection) in the presence of synaptic blockers and under gap-free conditions to allow continuous monitoring of spontaneous changes in membrane potential and firing activity before and after treatment with 200 nM semaglutide. All cells in which we injected a modest amount of current (5 pA, 1 second) were able to fire trains of action potentials (9/9 POMC-GFP and 8/8 POMC-NG). Despite the hyperpolarizing effect of synaptic blockers, about half of recorded cells (3/7 POMC-GFP and 4/8 POMC-NG) showed spontaneous action potentials in the absence of current injection, suggesting that hPSC-derived POMC neurons were healthy and capable of stimulus-evoked electrical activity.

Next, we selected cells that gave stable recordings for at least 10 minutes before and 20 minutes after administration of 200 nM semaglutide for 6 minutes in HBSS. We found that application of 200 nM semaglutide induced a robust depolarization and increased action potential firing that persisted upon drug withdrawal (Fig. 3A,B,F; Fig. S3; Table S3). Specifically, in POMC neurons derived from POMC-GFP and POMC-NG reporter lines, semaglutide significantly depolarized the membrane potential by approximately 15 mV (95% CI: 9 to 21) and 10 mV (95% CI: 5 to 14) respectively (Fig. 3A-E; Fig. S3; Table S3), and increased the action potential firing rate of POMC neurons by an average of 14 Hz (95% CI: 3 to 24 Hz) and 17 Hz (95% CI: -1 to 35 Hz), respectively (Fig. 3A-C, F,G; Fig. S3; Table S3). Next we performed instantaneous firing frequency analysis on neurons that were spontaneously active at baseline by calculating the reciprocal of the interspike interval (ISI) to give insight into neuronal firing dynamics (Fig. 3H). Semaglutide application induced significant and persistent shifts of the instantaneous frequency value distribution from the baseline (median: 0.33 Hz; IQR: 0.14 -0.94) to post-semaglutide exposure (median: 1.19 Hz; IQR: 0.79 -1.65). These results indicate that POMC neurons fired at progressively shorter ISIs during and after semaglutide exposure (Fig. 3I,J, Table S3). Together, electrophysiological and calcium imaging data provide convergent evidence that semaglutide activates hPSC-derived POMC neurons through post-synaptic and GLP-1R-dependent mechanisms to depolarize and excite hypothalamic POMC neurons and drive sustained calcium influx, likely through voltage-gated calcium channels (VGCC) and/or non-specific cation channels.

**Figure 3.**
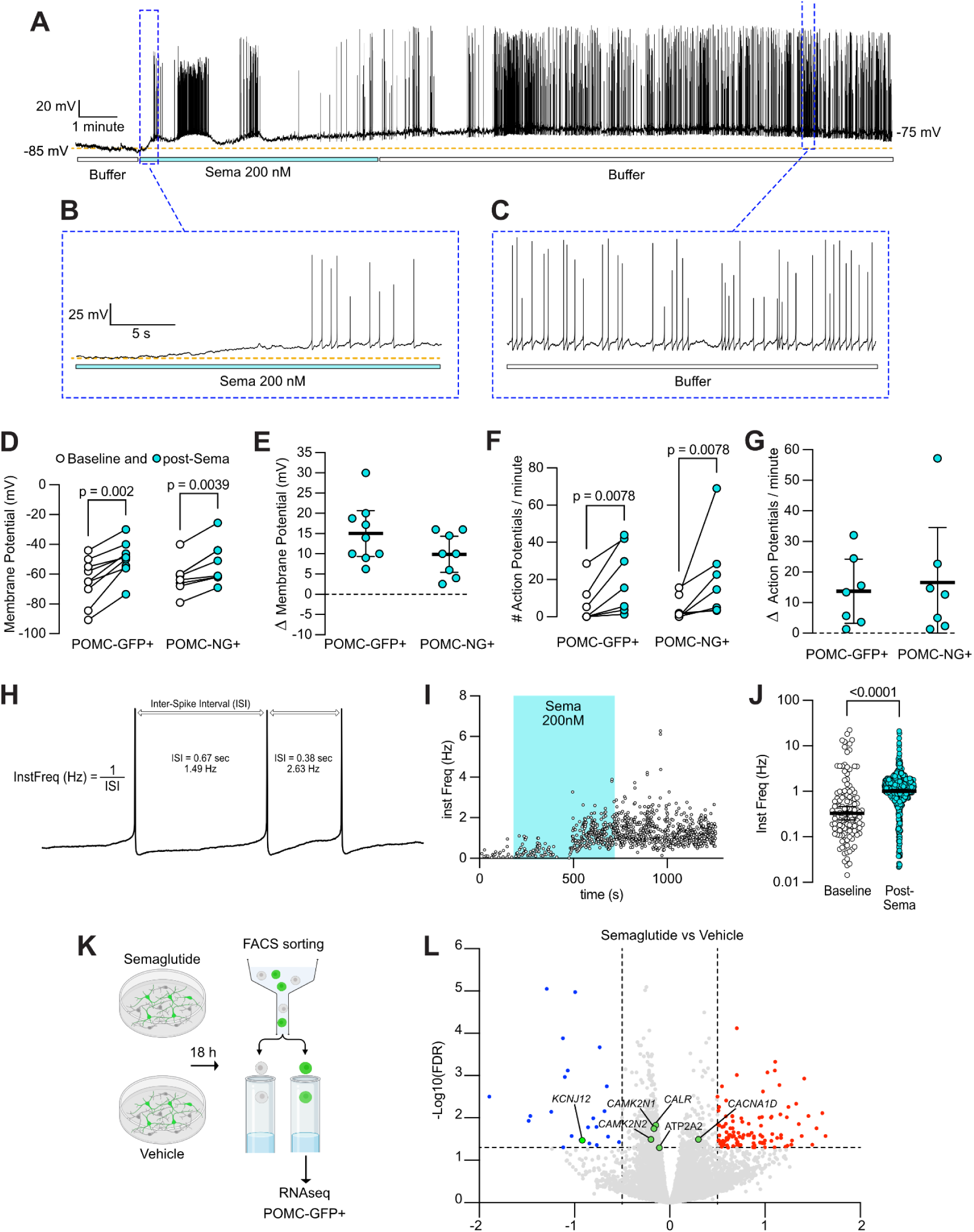
Electrophysiological and transcriptional responses of POMC neurons to semaglutide. **A-C)** Representative trace of a hPSC-derived POMC neuron under perforated patch preparation that is relatively quiescent under baseline conditions (A) but is depolarized upon administration of 200 nM semaglutide leading to an increased rate action potential firing (B) that persists for tens of minutes after semaglutide withdrawal (C). **D,E)** Semaglutide administration significantly depolarized the membrane potential of green fluorescent POMC neurons in two genetically distinct knock-in cell lines (POMC-GFP+ n=9; POMC-NG+ n=8) by an average of 15 mV (95% CI: 9 to 21 mV) and 10 mV (95% CI: 5.4 to 14.3 mV) mV respectively (E). **F,G)** Semaglutide administration significantly (p<0.01) increased (F) the action potential firing rate (F) by 14 Hz (95% CI: 3 to 24 Hz) and 17 Hz (95% CI: -1 to 35 Hz), respectively (G). **H-J)** The instantaneous frequency, defined as the reciprocal of the inter-spike interval, (H) increased upon 200 nM semaglutide administration as illustrated in a representative recording (I), resulting in a significantly shifted distribution (p<0.0001, unpaired Mann-Whitney t-test) of the instantaneous frequency values (**J**). Each dot represents an instantaneous frequency measurement from n=4 neurons (unpaired Mann-Whitney t-test). **K)** Schematic of hPSC-derived hypothalamic neurons treatment with vehicle or 1 𝛍M semaglutide for 18 hours followed by FACS purification of GFP+ cells **L**) Volcano plot showing significantly enriched (red) or depleted (blue) genes in semaglutide-treated vs. vehicle-treated neurons. Enriched or depleted genes of interest are highlighted by green dots. (n=6 technical replicates/group).

**Figure S3.**
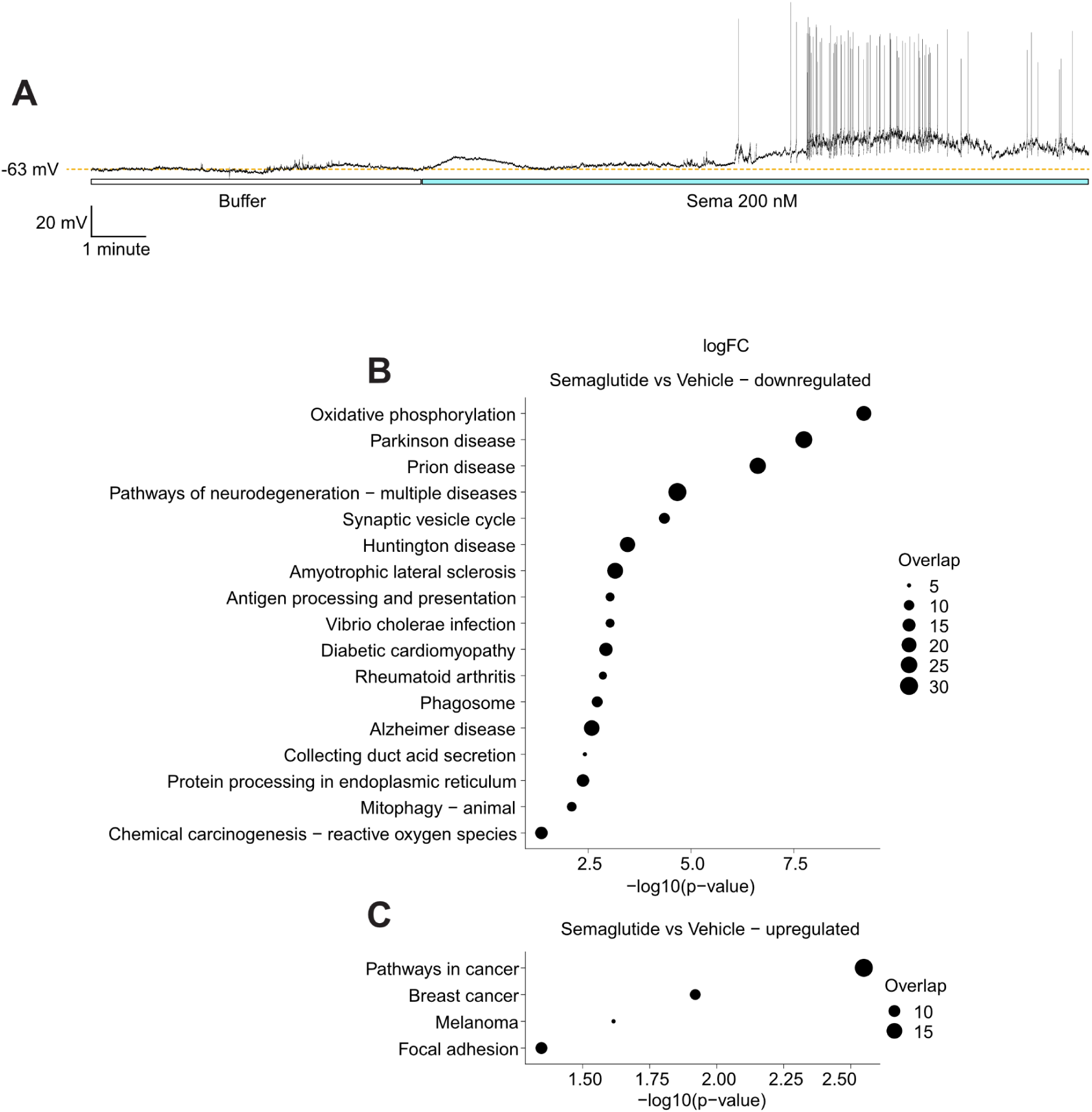
Electrophysiological transcriptional pathways altered by semaglutide in POMC neurons. **A)** Representative trace of a hPSC-derived POMC-NG+ neuron under perforated patch preparation that was silent under baseline conditions but depolarized after administration of 200 nM semaglutide, leading to an increased action potential firing rate and increased resting membrane potential. **B,C)** KEGG enrichment analysis of significantly (FDR<0.05) downregulated (**B**) or upregulated (**C**) genes.

### Semaglutide alters gene expression in POMC neurons

Since semaglutide likely drives the sustained activation of appetite-regulatory neurons *in vivo*, we wondered what impact it would have on the transcriptome of human POMC neurons *in vitro*. We therefore treated cultures of hypothalamic cells differentiated from *POMC-GFP* reporter hPSCs with vehicle or semaglutide for 18 hours (Fig. 3K), purified GFP+ cells by FACS from six replicate cultures per treatment group, and analyzed their transcriptomes by bulk RNA sequencing. After adjusting for multiple comparisons, we found that semaglutide significantly (FDR<0.05) down-regulated 393 genes and up-regulated 257 genes (Fig. 3L; Table S3C), of which 23 and 107 genes showed a log_2_(|fold-change|)>0.5, respectively. Pathway analysis (Kanehisa et al. 2023) of these differentially expressed genes revealed significant upregulation of pathways associated with cell survival and adhesion (Fig. S3C), and a significant downregulation of pathways associated with oxidative stress and neurodegeneration, among others (Fig. 3B).

Specifically, we observed significant changes in genes that may help sustain higher levels of neuronal activity including the enriched expression of the calcium channel *CACNA1D* (Fig 3L; Table S3C), which corresponds to an L-type VGCC targeted by the pharmacological blockers applied in this study (Fig 4D; Table S4B). Semaglutide treatment also significantly downregulated the inward-rectifying potassium channel *KCNJ12* (Kir2.2) (Fig S3L; Table S3C), with the predicted effect of increasing neuronal excitability since Kir2 channels contribute to the stabilization of the neuronal resting membrane potential and reduce the action potential threshold in hypothalamic (Smith et al. 2006) and other neurons (Hibino et al. 2010; Ford and Baccei 2016). In contrast, voltage-gated sodium channel genes (SCN family) and the sodium leak channel *NALCN* were unchanged. Multiple genes involved in intracellular Ca^2+^ handling and Ca^2+^-dependent signaling, including *ATP2A2*, *CALR*, and the CaMKII inhibitors *CAMK2N1* and *CAMK2N2*, were significantly downregulated (FDR<0.05) (Fig 3L; Table S3C), perhaps in response to prolonged increases in [Ca^2+^]_i_. We also observed modest but significant changes in the expression of genes associated with obesity, including *FAIM2* which encodes an anti-apoptotic protein associated with childhood obesity (Littleton et al. 2024) and *NTRK2* which encodes the receptor for BDNF (Yeo et al. 2004). Together, these data suggest that prolonged semaglutide exposure might remodel the excitability of POMC neurons, while preserving the machinery required for action potential initiation.

**Figure 4.**
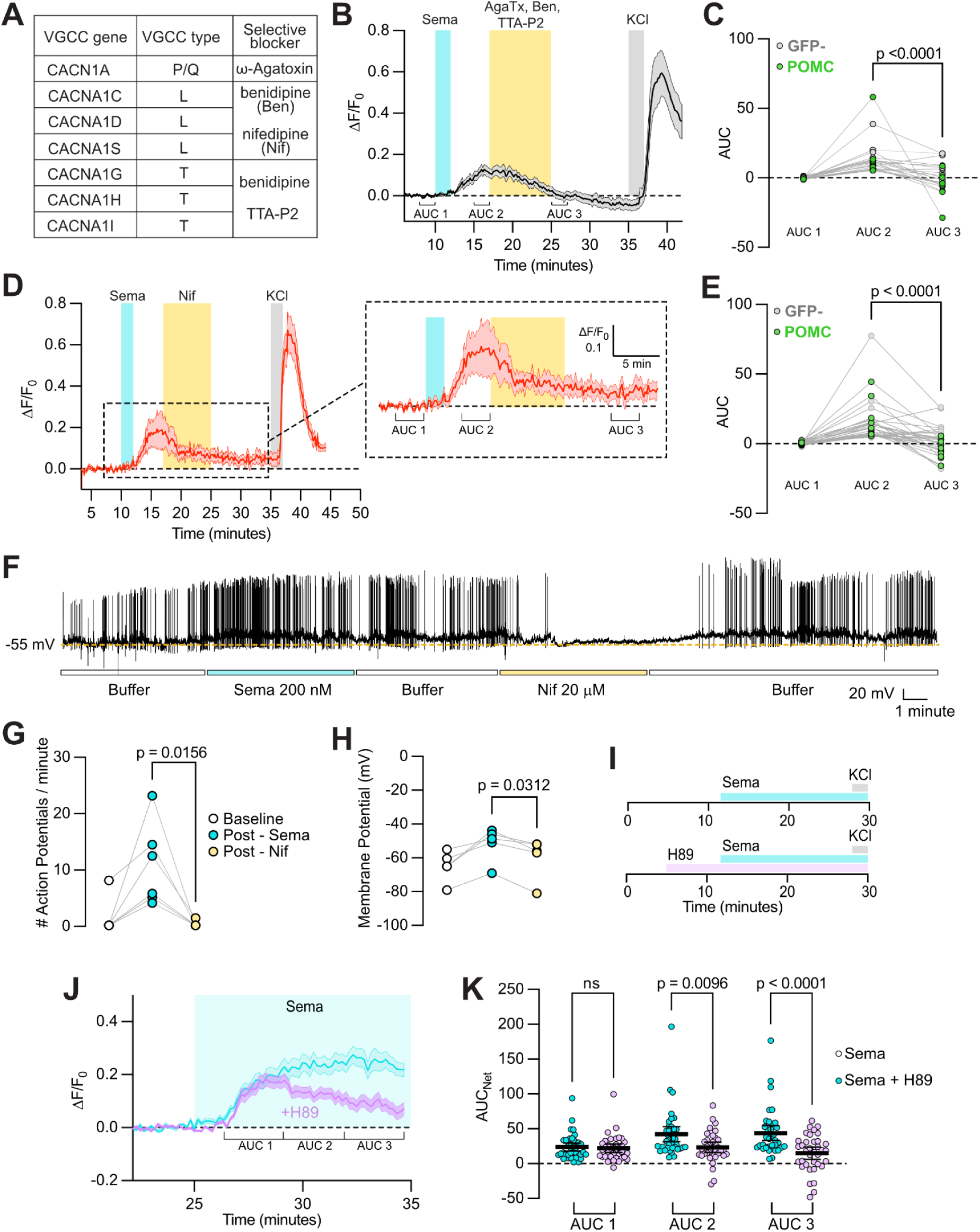
Durable responses to GLP-1R agonists require L-type voltage-gated calcium channels and PKA. **A)** Table of voltage-gated calcium channels (VGCC), channel types, and selective blockers of these channels. **B-E)** Co-application of a cocktail of VGCC blockers (B, yellow) significantly (p<0.0001; paired nonparametric Wilcoxon t-test) reversed semaglutide-induced calcium indicator fluorescence (C), and these results were recapitulated by L-type selective VGCC blocker nifedipine (Nif, D) in both GFP-neurons and POMC-GFP+ neurons (E). **F)** Electrophysiological recording trace of a POMC neuron showing its depolarization and increased action potential firing rate upon stimulation by 200 nM semaglutide (blue) that persisted after washout with buffer (white) but was reversed by nifedipine (yellow). **G,H)** The effects of L-type VGCC blockade on action potential firing rate (G) and membrane potential (H) in semaglutide-simulated POMC neurons are statistically significant and reproducible (n=6 and n=5, respectively). **I-K)** Administration of the PKA inhibitor H89 (pink) prior to the administration of semaglutide (blue; **I**) does not alter the early calcium response of cells but significantly reduces (p<0.01, Mann-Whitney t-test) sustained calcium indicator fluorescence as shown in traces of average cell responses ± SEM (J) and as quantified by taking the area under the curve in successive time windows (K), where lines indicate mean and 95% CI, and dots indicate n=39 semaglutide-treated neurons and n=36 semaglutide + H89 treated neurons.

**Figure S4.**
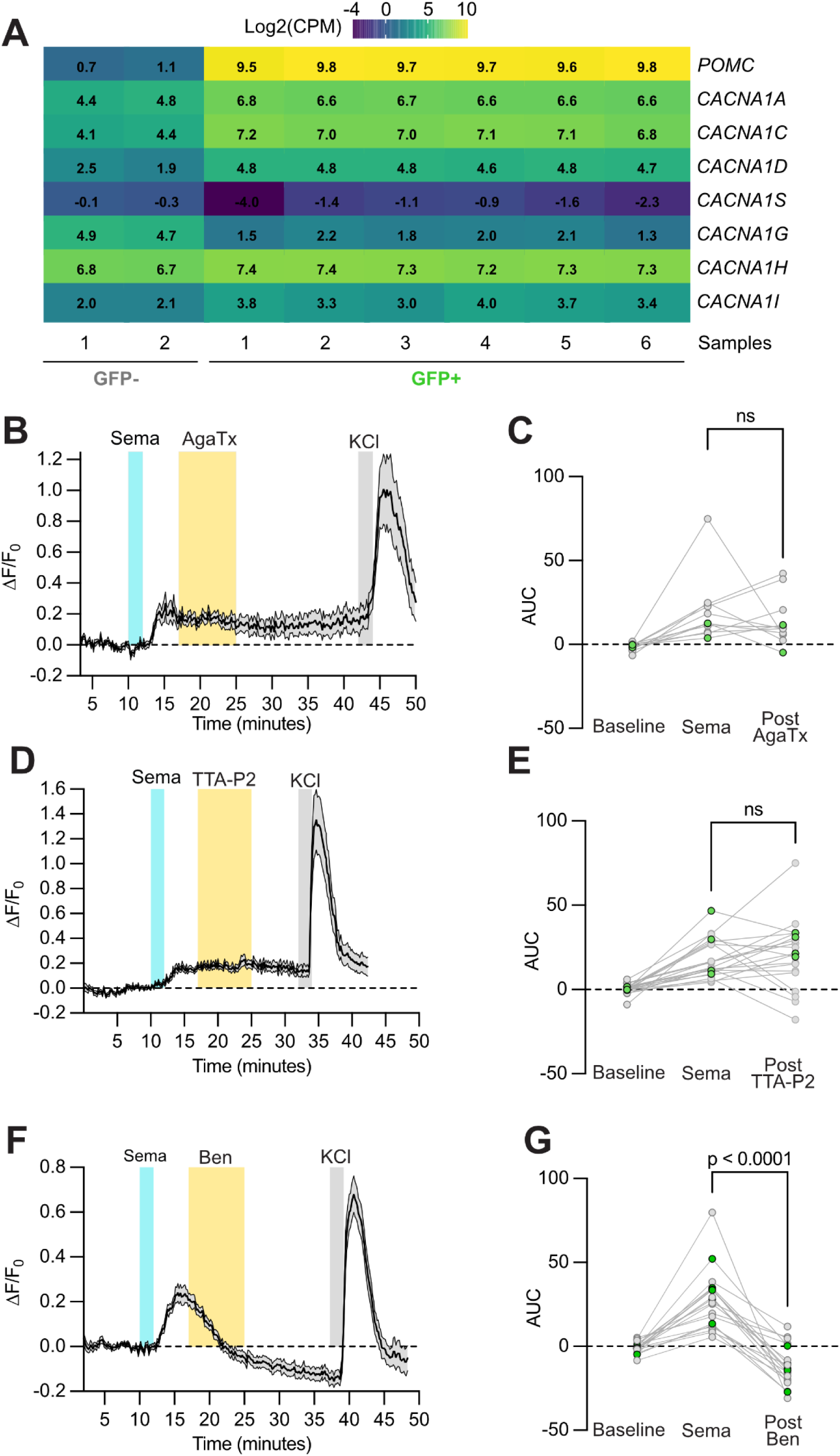
GLP-1R agonist-induced calcium responses require voltage-gated calcium channels. **A)** Expression of VGCCs from transcriptomic analysis of FACS-purified GFP- and GFP+ (POMC) neuron populations. **B-G)** Administration of the P/Q-type VGCC blocker ω-Agatoxin (B,C; n=11 cells) or the T-type VGCC blocker TTA-P2 (D,E; n=20 cells) did not significantly decrease semaglutide-induced calcium indicator fluorescence, as seen average traces (B,D) and in a quantification of GFP- and POMC-GFP+ cells (C,E). **F, G)** Panels as in C-E but with the L- and T-type VGCC blocker benidipine, which significantly reduced (p<0.0001, paired nonparametric Wilcoxon t-test) semaglutide-induced calcium dye fluorescence (n=19 cells).-

### Neuronal responses to semaglutide require L-type calcium channels and PKA

We hypothesized that the persistent electrical activation and increased [Ca^2+^]_i_ of human POMC neurons in response to semaglutide requires VGCCs as described in pancreatic β-cells, where L-type calcium channels are engaged upon GLP-1R agonist binding (Gomez et al. 2002). We therefore examined bulk RNAseq data from FACS purified hPSC-derived GFP+ and GFP-cells, and found that most calcium channels we tested were expressed in POMC neurons, including the L-type VGCCs *CACNA1C and CACNA1D* (Fig. S4A).

To determine whether VGCCs contribute to the prolonged increase in [Ca^2+^]_i_ we observed, we first took baseline measurements (AUC 1) and then treated cultures with 200 nM semaglutide for two minutes to identify activated cells, then waited 5 minutes for responses to plateau, and finally added a cocktail of potent small molecule VGCC blockers for 8 minutes (Fig. 4A) including 10 μM benidipine, 30 nM TTA-P2, and 30 nM ω-Agatoxin (Zamponi et al. 2015). We found that upon adding these channel blockers, the semaglutide-induced calcium signal (AUC 2, 2 minutes after semaglutide withdrawal) dropped significantly (p<0.05) for both POMC and GFP-neurons after blocker treatment and withdrawal (AUC 3, Fig. 4C, Table S4). These findings suggest that the persistent semaglutide-induced increase in [Ca^2+^]_i_ requires VGCCs.

To isolate which channel blockers drove this effect, we repeated these studies with one candidate VGCC blocker at a time. We saw no significant drop in calcium indicator fluorescence intensity in semaglutide-activated cells upon administration of either 30 nM ω-Agatoxin (Fig. S4B,C; Table S4) or 30 nM TTA-P2 (Fig. S4D,E; Table S4), but observed a significant (p<0.01) drop in fluorescence intensity in response to 10 μM of the high affinity L-type VGCC blocker benidipine (Fig. 4F,G, Table S4). To confirm these findings, we replicated these experiments with the structurally distinct L-type VGCC blocker nifedipine and again observed a significant (p<0.0001) drop in fluorescence intensity (Fig. 4D,E; Table S4). Together, these findings suggest that L-type VGCCs are required for the sustained increase in [Ca^2+^]_i_ observed in human POMC neurons.

To confirm this interpretation, we performed perforated patch-clamp recordings on hPSC-derived GFP⁺ and NG⁺ POMC neurons as previously described. At baseline (pre-semaglutide administration), neurons displayed an average membrane potential of -64.9 mV (95% CI: -75.94 to -53.86) and an average action potential firing rate of 1.3 Hz (95% CI: -2.1 to 4.8 Hz). Treatment of cultures with 200 nM semaglutide significantly depolarized the membrane potential to an average of -51.7 mV (95% CI: -64.2 to -39.2 mV) and increased the action potential firing rate to 9.4 Hz (95% CI: 1.7 to 17.0 Hz) (Table S4A, p<0.05), even after washout with HBSS (Fig. 4F-H). Application of 20 μM nifedipine returned both the firing rate (0.22 Hz; 95% CI: -0.35 to 0.79) and membrane potential (−59.6 mV; 95% CI: -74.72 to -44.48 mV) of POMC neurons to levels indistinguishable (p>0.05) from baseline (Fig S4A). Quantification across multiple recordings showed the effects of L-type VGCC blockade on action potential frequency (Fig. 4G; Table S4) and membrane potential (Fig. 4H; Table S4) were both significant (p<0.05) and reproducible. Together, the data show that L-type VGCCs likely maintain semaglutide-induced excitability in human POMC neurons.

We wondered how GLP-1R activation is mechanistically linked to increased calcium influx through L-type VGCCs. In cardiomyocytes and other neuron types, GPCRs can directly modulate that activity of L-type calcium channels (Dolphin, 2018). Specifically, Gα_s_-coupled receptors (e.g. β-adrenergic receptors) stimulate adenylate cyclase to increase intracellular cAMP concentration, which activates protein kinase A (PKA) leading to the phosphorylation of serine residues on the C-terminal tail of L-type VGCCs (Mitterdorfer et al., 1996) to increase the likelihood of channel opening (Cachelin et al., 1983), thereby increasing inward Ca^2+^ currents (Catterall 2011; Budde et al. 2002). To test the hypothesis that GLP-1R activation in POMC neurons requires PKA to mediate durable depolarization, we performed calcium imaging on hPSC-derived hypothalamic neurons in the presence or absence of 10 μM H-89, a PKA inhibitor, prior to semaglutide application (200 nM). We found that PKA inhibition did not affect the timing or magnitude of the initial calcium response, but markedly reduced the sustained intracellular [Ca^2+^]_i_ increase induced by semaglutide (Fig. 4I-J). Quantification of the AUCs across successive time windows (Fig. 4K) confirmed progressive and significant reductions (AUC 1 p>0.05; AUC 2, p<0.01; AUC 3, p<0.0001) in the prolonged calcium signal over time in semaglutide and H-89-treated neurons (n=36) compared with neurons treated with semaglutide alone (n=39). These results indicate that PKA activity is required to sustain semaglutide-induced calcium elevations in human POMC neurons. We conclude that the prolonged response of appetite-regulatory human neurons to GLP-1R agonists is likely mediated by PKA-dependent action on L-type VGCCs (Fig. 5).

**Figure 5.**
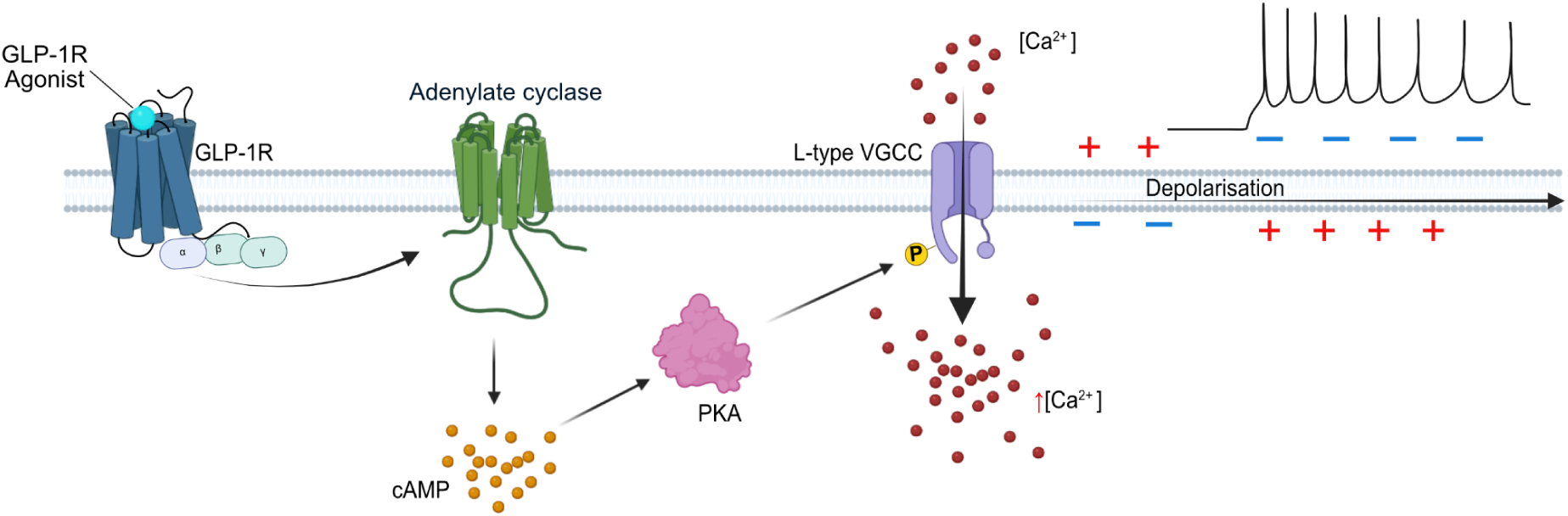
Working model of GLP-1R-mediated excitation of human hypothalamic neurons. Experimental data collected in this study are consistent with a model in which GLP-1R activation by an agonist leads to Gα_s_-mediated activation of an adenylate cyclase, leading to an increase in intracellular cAMP concentration that stimulates protein kinase A (PKA) that in turn phosphorylates an L-type voltage-gated calcium channel to increase its conductance, contributing to sustained neuronal depolarization and increased firing rate.

## Discussion

The remarkable success of GLP-1R agonist drugs in humans follows a long history of research in animal models, where candidate mechanistic targets in the hindbrain, hypothalamus, and other brain regions have been identified (Müller et al. 2019). While histological and single-cell studies confirm that *GLP1R* is expressed in the human hypothalamus (Tadross et al. 2025), the accessibility of these cells has previously precluded their functional analysis. In this study, we used complementary methodologies across two genetically distinct reporter cell lines to establish hPSC-derived hypothalamic neurons as a model for investigating human neuronal GLP-1R signaling. Below, we discuss the implications, outstanding questions, and the strengths and limitations of our study.

A key first step in our investigation was to determine whether human iPSC-derived POMC neurons recapitulate the *GLP1R* mRNA expression patterns observed in the rodent (Steuernagel et al. 2022) and human hypothalamus (Tadross et al. 2025). Consistent with the literature, our data from single-cell and bulk transcriptomics and RNAscope revealed that *GLP1R* transcripts are present in many hypothalamic neurons and significantly enriched in POMC-expressing cells. Next, using calcium imaging we observed that the majority of hPSC-derived POMC neurons responded to GLP-1R agonists. We observed that even a brief exposure of human POMC neurons to GLP-1R agonists resulted in a prolonged (>20 minutes) depolarization of their membrane potential, increased action potential firing rate, and elevation of [Ca²⁺]ᵢ. These findings are consistent with previous studies in rodent POMC neurons that also showed long-lasting increased excitability in response to GLP-1R agonists (Dong et al. 2021; Secher et al. 2014). In future studies, it would be interesting to determine the full duration of this increased activity, the kinetics by which [Ca^2+^]_i_ returns to baseline, and compare these kinetics to any changes in intracellular cAMP concentrations to study responses more proximal to putative GLP-1R-Gα_s_ coupling.

Electrophysiological analysis and pharmacological blockade of candidate channels suggested that the durable excitability we observed is likely mediated by PKA-dependent action on L-type VGCCs. We found that while the PKA inhibitor H-89 markedly reduced the sustained calcium signal, it did not appear to suppress the initial calcium response of POMC neurons, suggesting the involvement of a separate and still unidentified pathway to increase [Ca²⁺]ᵢ. While our pharmacological approach could not distinguish between different L-type VGCCs, we speculate that CACNA1D may mediate the responses we observed, since this channel is expressed at appreciable levels in hPSC-derived POMC neurons (∼28 CPM) and is transcriptionally induced by semaglutide (Fig 3L; Table S3C). However, we cannot exclude the involvement of other channels, such as non-selective cation channels thought to mediate the exendin-4-induced depolarization of hypothalamic hypocretin neurons (Acuna-Goycolea and van den Pol 2004). Treatment with the L-type selective VGCC blocker nifedipine suppressed the signal of fluorescent calcium indicators to baseline levels (Fig. 4D,E) and this effect was even more pronounced with benidipine or a cocktail of VGCC blockers (Fig. 4B, S4F). The suppression of [Ca²⁺]ᵢ to levels below baseline may be due in part to the contribution of blockade of open VGCCs on neurons that activate near neuronal resting membrane potential (∼-65 mV) (Perez-Reyes 2003; Huguenard 1996), and/or non-specific effects of benidipine.

At the transcriptional level, we observed that prolonged semaglutide exposure changed the expression of genes associated with the maintenance of neuronal excitability and pathways related to intracellular Ca²⁺ handling and Ca²⁺-dependent signaling in neurons. These genes included *ATP2A2* which encodes the ER Ca²⁺ pump *SERCA2* that mediates cytosolic Ca²⁺ sequestration into the endoplasmic reticulum (Nakajima et al. 2021), the ER luminal Ca²⁺-binding chaperone *CALR* whose product contributes to intracellular Ca²⁺ buffering and signaling regulation (Coe and Michalak 2009), and the CaMKII inhibitory genes *CAMK2N1* and *CAMK2N2* which encode endogenous suppressors of CaMKII activity that modulate Ca²⁺-dependent signaling pathways in neurons (Nelson et al. 2005). Together, these changes suggest that prolonged semaglutide administration altered ER Ca²⁺ buffering and remodeling of Ca²⁺-dependent signaling. Additionally, we found that semaglutide treatment was associated with the significant downregulation of pathways associated with neurodegeneration. While these findings require validation and mechanistic follow-up, it has not escaped our attention that GLP-1R agonists show neuroprotective and cognitive benefits to mice (During et al. 2003; Li et al. 2009; Panagaki et al. 2018), and humans (Wu et al. 2018; Vadini et al. 2020). While the mechanisms by which GLP-1R agonists exert their neuroprotective effects remain largely obscure (Reich and Hölscher 2022; Kopp et al. 2022), our findings suggest that they could involve direct effects on neurons. Overall, our findings are broadly consistent with reports from previous studies in rodents (Rønnekleiv et al. 2014; He et al. 2019; Péterfi et al. 2021; Secher et al. 2014), establishing hPSC-derived hypothalamic neurons as an evolutionarily conserved, disease-relevant, functional, and scalable human model system for studying GLP-1R biology.

### Limitations of study

Though we took care to ensure the reproducibility and relevance of our work, we acknowledge several limitations. Specifically, while we analyzed hPSC-derived neurons at time points when they fired spontaneous action potentials and showed clear calcium responses, they are unlikely to show the same functional responses as mature primary neurons. Furthermore, while we added a cocktail of synaptic blockers in all experiments, we cannot formally exclude the possibility that cells communicated with each other via paracrine or other signals. We did not consider the contributions of non-neuronal cells in hypothalamic cultures, or assess non-hypothalamic but GLP-1R-responsive cell types of interest, such as neurons of the dorsal vagal complex.

The identity of GFP-cells that responded to GLP-1R agonists is unknown, other than that they were likely neuronal based on their morphology and robust responsiveness to KCl. We found that a small but reproducible fraction of these GFP-neurons appeared to reduce intracellular Ca²⁺ levels upon administration of GLP-1R agonists, as suggested by a sustained decrease in calcium indicator dye fluorescence. To our knowledge, evidence that GLP-1R agonists are able to reduce directly the intracellular Ca²⁺ levels has not been reported although evidence for indirect inhibition has been described for Agouti-related protein (AgRP) neurons from *ex vivo* slice recordings (Dong et al. 2021; Secher et al. 2014).

Much attention has been paid to the heterogeneity of POMC neurons based on the genes they express, their anatomical location and connectivity, or their functional responses to factors like leptin and GLP-1 as determined by electrophysiological recording or calcium imaging (Quarta et al. 2021). Since sub-populations can be defined in many different ways, there is no consensus about their precise composition, or how directly mouse sub-populations might map onto hPSC-derived or primary human counterparts. To functionally define hPSC-derived human POMC neuron heterogeneity requires characterization beyond what is reported in this study, including but not limited to cAMP and calcium imaging in response to a broad panel of candidate ligands, followed by confirmation in primary human brain tissue. The findings presented here constitute an important step in this direction.

## Materials and methods

### Human pluripotent stem cell (hPSC) culture and hypothalamic differentiation

All cultures were maintained at 37°C in humidified incubators at 5% CO_2_ and 20% O_2_. Human pluripotent stem cell (hPSC) line HUES9-POMC-GFP (passage 12-18 after reporter generation) and KOLF2.1J-POMC-NLS-Neongreen (POMC-NG; passage 21-26, passage 5-10 after reporter generation) were generated (Chen, Yang, et al. 2023) and maintained as previously described (Chen, Mazzaferro, et al. 2023). Briefly, stem cells were maintained on Geltrex-coated plates in StemFlex medium and passaged upon reaching ∼70% confluence with 0.5 mM EDTA in the presence of 10 µg/mL Rock Inhibitor Y-27632.

These cell lines were then differentiated into hypothalamic neurons as previously described (Chen, Mazzaferro, et al. 2023; Merkle et al. 2015; Kirwan et al. 2017). Briefly, hPSCs were dissociated to a single-cell suspension and plated onto Geltrex-coated plates at a density of 1 x 10^5^ cells/cm^2^ in StemFlex medium supplemented with 10 µg/mL Rock Inhibitor Y-27632. The following day, media was changed to N2B27 media containing small molecule BMP, TGFβ and WNT inhibitors to promote neural induction, followed by small molecule SHH agonists to promote ventralization. On day 13 of differentiation, hypothalamic progenitors were dissociated with a mixture of TrypLE Express and 200 U/mL papain, washed with media supplemented with 2 mg/mL DNAse I (1:60), and re-plated at a concentration of 2.5 x 10^5^ cells/cm^2^ onto plates coated with 4 μg/mL iMatrix-511 (Takara Bio) in N2B27 medium supplemented with 10 μg/mL BDNF and 10 µg/mL Rock Inhibitor Y-27632. The following day, media was changed to Synaptojuice medium 1 (N2B27 medium with 2 μM PD0332991 isethionate, 5 μM DAPT, 370 μM CaCl_2_, 1 μM LM22A4, 2 μM CHIR99021, 300 μM GABA, and 10 μM NKH447) supplemented with 10 μg/mL BDNF. After 1 week, media was switched to Synaptojuice 2 (N2B27 medium with 2 μM PD0332991 isethionate, 370 μM CaCl_2_, 1 μM LM22A4, and 2 μM CHIR99021) and supplemented with 10 μg/mL BDNF (Kemp et al. 2016). Cultures were maintained by media change every two days.

### RNA Scope

Hypothalamic cultures (HUES-POMC-GFP and KOLF2.1J-POMC-NG) at 40 days post-differentiation were dissociated, washed with Dulbecco’s Phosphate-Buffered Saline (DPBS^-/-^), and approximately 10 million cells per cell line were fixed in 10% neutral buffered formalin for 4 hours at room temperature on a rotating mixer. Cells were then centrifuged at 500 x g for 3 minutes and decanted the fixative. The cell pellets were resuspended with DPBS^-/-^ with 1% bovine serum albumin (BSA) and centrifuged at 500 x g for 3 minutes. The resulting pellets were amalgamated in liquid HistoGel™ and embedded into a paraffin block using a standard embedding procedure.

Simultaneous detection of *GLP1R* and *POMC* was performed on cell pellet sections using the RNAscope® 2.5 LS Multiplex Reagent Kit, RNAscope® 2.5 LS Probe GLP1R-CDS, and RNAscope® 2.5 LS Probe POMC (ACD, Hayward, CA, USA). Briefly, sections were cut at 3 µm thick, incubated for 1 hour at 60°C before loading onto a Bond RX instrument (Leica Biosystems). Slides were deparaffinized and rehydrated on board prior to pre-treatments using Epitope Retrieval Solution 2 (Leica Biosystems) at 88°C for 15 minutes, and ACD Enzyme from the Multiplex Reagent kit at 40°C for 15 minutes. Probe hybridization and signal amplification was performed according to the manufacturer’s instructions. Probes were visualized using Opal fluorophores (Opal 570 for GLP1R and Opal 520 for POMC; Akoya Biosciences) diluted to 1:500 and 1:1000, respectively, using RNAscope LS Multiplex TSA Buffer. Slides were then removed from the Bond Rx and mounted using ProLong™ Diamond Antifade Mountant (Thermo Fisher Scientific).

The slides were then imaged on the Akoya PhenoImager HT to create whole slide scans. Images were captured at 40x magnification with a resolution of 0.25 μm per pixel and were analyzed with HALO Image Analysis Software v4.0.5107.357 (Indica Labs Inc., USA). *GLP1R* and *POMC* positive cells were detected in the HALO®FISH module based on intensity thresholds set for each cell line using negative controls. Nuclei were identified by DAPI staining, and the cytoplasm was extrapolated 3 μm around the nucleus. The accuracy of cell segmentation and the detection of *GLP1R* and *POMC* transcripts were verified visually to confirm the absence of false negatives. The number of spots per cell in the *GLP1R* channel was then quantified automatically.

### Analysis of single-cell sequencing data

Processed single-cell RNA sequencing data obtained from genetically distinct hPSC lines differentiated to hypothalamic neurons (Chen, Yang, et al. 2023) as well as published data from HypoMap mouse single-cell hypothalamus atlas (Steuernagel et al. 2022) were analyzed for the expression of *GLP1R* and other genes of interest using expression cutoffs of ≥1 UMI.

### Preparation of drugs and chemicals

Peptides and small molecule drugs used in this study were reconstituted according to manufacturer instructions, aliquoted into single-use vials, and stored at -70°C or -20°C until the day of each experiment.

### Calcium imaging

Calcium Imaging experiments were performed as previously described (Chen, Mazzaferro, et al. 2023). Briefly, hypothalamic progenitors differentiated from the HUES9-POMC-GFP cell line were replated onto 35 mm x 10 mm polystyrene imaging dishes (Corning) coated with 1 µg/cm^2^ of iMatrix-511. All analyses were performed on neurons 40-50 days post-differentiation and transitioned from SynaptoJuice medium to BrainPhys medium 24-48 hours prior to recording. On the day of recording, neuronal cultures were loaded with Cal-590 AM fluorescent calcium indicator dye (Stratech) following manufacturers’ instructions. An extracellular bath solution consisting of Hanks’ Balanced Salt Solution without Phenol Red (HBSS; ThermoFisher Scientific) and synaptic blockers were used for all aspects of calcium imaging, including dye loading, perfusion, and dilution of GLP-1R agonists and other experimental substances. Synaptic blockers were added to final concentrations of 100 µM DL-AP5, 50 µM Picrotoxin, 30 µM CNQX, and 20 µM Strychnine to inhibit NDMA, GABA, AMPA/Kainate, and glycine receptors, respectively. Immediately before each experiment, GLP-1R agonists were diluted from stock solutions to final concentrations of 200 nM in unless otherwise indicated.

Imaging dishes were placed on an Olympus BX51WI Fixed Stage Upright Microscope and imaged using a 16-bit high-speed ORCA Flash4.0 LT plus digital sCMOS camera. Neurons were identified by Cal-590 AM fluorescence using an excitation wavelength of 540 nm and an emission wavelength of 590 nm. Images were taken using a 20x objective and acquired at a frequency of 0.1 Hz (100 ms exposure/frame) using a CoolLED pE-300 white illumination system and HCImage software for acquisition. Neurons were perfused continuously at room temperature (3 mL/min) using a gravity-driven perfusion system for the entire length of the experiment. For recordings, cells were perfused with extracellular bath solution for 10 minutes to record baseline fluorescence, and then perfused with extracellular bath solution containing the experimental compound for two minutes, followed by a second washout period (10-20 minutes) and stimulation with 50 mM KCl for two minutes followed by a washout period to identify responsive neurons. Videos of fluorescence intensity in the green (GFP) and red (Cal-590 AM) channels were recorded at an acquisition rate of 6 frames per minute.

For some experiments with exendin-(9-39), calcium imaging was performed in 96-well plates on an Opera Phenix (Revvity) high content microscope. Hypothalamic cultures were re-plated at approximately 1.5×10^5^ cells/cm^2^, maintained as described above, and moved to BrainPhys medium lacking phenol red the day before imaging. After loading cells with calcium dye, counter-staining with Hoechst, and adding synaptic blockers as described above, cultures were imaged at 37.5°C and 5% CO₂ in confocal mode with a 20× water immersion objective and 2x binning. Four fields of view were acquired from each well, and a maximum of 30 wells was imaged per experiment to ensure that the time between sequential acquisitions of the same field did not exceed 154 seconds. Images were captured in the Hoechst (405 nm excitation and 435–480 nm emission), GFP (88 nm excitation and 500-550 nm emission), and calcium dye (561 nm excitation and 570-630 nm emission) channels at each timepoint. Fluorescence measurements were acquired at seven baseline timepoints, after which 10 μL of ligand or vehicle solution (prepared as a 10× the desired final concentration, dissolved in BrainPhys-based preparation medium) was gently added to each well, which already contained 90 μL of imaging medium. After capturing an additional seven time points, 10 μL of 0.5 M potassium chloride (KCl) solution was then added per well to achieve a final concentration of approximately 45 mM KCl.

### Imaging and figure preparation

Images of hypothalamic cultures were collected on the calcium imaging microscope as described above, and separate cultures were plated onto PhenoPlateTM 96-well microplates (Perkin Elmer), fixed in 4% paraformaldehyde (Thermo Fisher Scientific) for 10 minutes, rinsed three times with Tris-buffered saline containing 0.1% Triton X-100 (TBS-T; Sigma-Aldrich), and incubated with primary antibodies including anti-α-MSH (A1H5, detects epitopes in the α-MSH region of POMC and developed by Professor Anne White; 1:5000) diluted in TBS-T containing 1% normal donkey serum (NDS; Stratech) overnight at 4°C. Cells were washed three times with TBS-T and incubated with secondary antibodies diluted in TBS-T with 1% NDS for 2 hours at room temperature on an orbital shaker. After the incubation with the secondary antibodies, cells were washed three more times with TBS-T.

Image acquisition was performed using the Perkin Elmer Opera Phenix Plus High-Content Screening System with a 20x or 40x water objective. Some figure elements were prepared with the help of BioRender, and some photomicrographs were background-subtracted, adjusted for brightness and contrast, and false-colored to enable merging of channels and aid in visualization. Figures were prepared in Adobe Illustrator or Affinity Designer.

### Quantification of responses from calcium imaging

To quantify changes in fluorescence intensity in GFP+ and GFP-neurons, regions of interest were manually drawn around candidate cells and raw fluorescence values from the 590 nm channel were exported to Microsoft Excel for each fluorescence time course. Only cells displaying a stable baseline prior to experimental compound administration and also a robust response to 50 mM KCl were considered for analysis. The change in fluorescence intensity as a function of time was expressed as (F − F_0_)/F_0_ or ΔF/F_0_, where F was the measured fluorescence intensity and F_0_ was the mean fluorescence intensity recorded during the ten-minute baseline period. Cellular responses to experimental compounds (e.g. GLP-1R agonists) were quantified by comparing the change in the area under the curve (AUC) of normalized fluorescence (ΔF/F_0_) intensity in a three-minute time window before and after addition of experimental compounds.

### Electrophysiology

Whole-cell recordings were obtained using the perforated Patch-Clamp technique as previously described (Goldspink et al. 2020). Briefly, microelectrodes were pulled from borosilicate glass (1.5 mm OD, 1.16 mm ID; Harvard Apparatus), and the tips were coated with refined yellow beeswax. Microelectrodes were fire-polished using a microforge (Narishige) and had resistances of 2-3 MΩ when filled with internal pipette solution. The internal pipette solution contained: 76 mM K_2_SO_4_, 10 mM NaCl, 10 mM KCl, 10 mM HEPES, 55 mM sucrose, and 1 mM MgCl_2_; adjusted to pH 7.2 with KOH. Amphotericin B (10 μg/ml) dissolved in DMSO was added to the pipette solution on the day of recording to perforate cell membranes. A silver/AgCl wire connected to the bath solution via a 0.15 M NaCl agar bridge was used as a ground.

Once cells had been patched in this configuration (Lippiat 2008), recordings were acquired using an Axopatch 200B amplifier connected through a Digidata 1440A A/D converter and pCLAMP software (Molecular Devices). Spontaneous action potential firing was recorded in current-clamp mode without injecting current (I=0). Neurons were perfused continuously at room temperature with HBSS solution (3 mL/min) in the presence of synaptic blockers using a gravity-driven perfusion system. For each experiment, we established a baseline recording period under perfusion, followed by perfusion with HBSS containing 200 nM semaglutide. Membrane potential and the spontaneous action potential firing were recorded for 10-30 minutes before and after applying semaglutide for six minutes. For each cell, mean membrane potential and action potential frequency were measured during 180-seconds-long time windows taken immediately before semaglutide application, and 60-120 seconds after semaglutide application, at which point semaglutide induced depolarization had plateaued.

### Semaglutide simulation and FACS purification of POMC neurons

To test for the transcriptional consequences of longer-term exposure to GLP-1R agonists, HUES9-POMC-GFP hypothalamic cultures at 40 ± 2 days post-differentiation were cultured in N2B27 medium for 72 hours before treatment with vehicle (DMSO) or semaglutide (1 µM, MedChem Express) in N2B27 medium for 18 hours. Cultures were then digested with TrypLE Express with 200 U/mL of papain and dissociated in wash media (N2B27) supplemented with 2 mg/mL DNAse I, 10 µg/mL Rock inhibitor, and 40 µg/mL actinomycin D. The cell suspension was centrifuged at 300 x g for 5 min and washed with wash medium and centrifuged again. The resulting cell pellets were resuspended with 500 µL of sorting buffer (DPBS^-/-^ with 1 mM EDTA, 25 mM HEPES, and 0.2% BSA) and transferred to DNA Lo-bind tubes for FACS sorting. Immediately before sorting, cells were filtered through a 40 µm Flowmi cell strainer and loaded onto the Aria Fusion Spyro set at 4°C and sorted using a 100 µm sort nozzle based on DAPI (0.1 µg/mL) for live/dead cells and GFP fluorescence. Sorted live cells were collected directly into RLT lysis buffer and RNA was immediately extracted using the RNeasy Plus Micro kit (Qiagen).

### Bulk RNA sequencing and analysis

RNA from sorted cells were analyzed on the Agilent 2100 Bioanalyser system using the Eukaryote Total RNA Pico kit (Agilent Technologies) and the mean RIN number was 9.5. Libraries of cDNA were then generated using the SMARTer Stranded Total RNA-Seq Kit v3 - Pico Input Mammalian (Takara). The resulting cDNA libraries were then quantified by the High Sensitivity DNA assay, pooled, and sequenced to a targeted depth of 50M reads/sample on Illumina flow cells to generate 100bp paired-end reads.

The resulting RNA sequencing libraries were pre-processed with TrimGalore (v0.6.7) (Krueger et al. 2021) to remove adapters and filter low-quality reads before aligning them to the human (GRCh38, Ensembl release 98) reference genome using STAR (v2.7.9a) (Dobin et al. 2013). Read counts per gene were then quantified using featureCounts (v2.0.0) (Liao et al. 2014) against the human (GRCh38, gencode v32) annotation. Qualimap (v2.2.2d) (Okonechnikov et al. 2016) was used to generate QC metrics of aligned reads for evaluating sample library quality, together with QC metrics produced by STAR and featureCounts. Size factor normalization and removal of lowly expressed genes were performed using edgeR (Robinson, McCarthy, and Smyth 2010). Further quality control analysis of the samples were performed using principal component analysis and expression correlation analysis to identify any outlier samples. For differential expression analysis, estimation of both negative binomial dispersions and quasi-likelihood dispersion was first performed using the edgeR package, followed by a quasi-likelihood F-test to identify genes which are differentially expressed between different treatment groups, with significance threshold set to FDR<0.05. Finally, gene set enrichment analysis of the differentially expressed genes was performed using the gprofiler2 R package (Raudvere et al. 2019), with custom background set to expressed gene list, analytical-adjusted p-value threshold set to 0.05 and data sources set to KEGG.

### Statistical analysis

Graphing and statistical analysis were performed using GraphPad Prism (version 7.02). For all one-way analyses of variance (ANOVA), the normality of residuals was first confirmed using the D’agostino Pearson test. For all instances in which multiple paired t-tests were used, data were first analyzed for normality using the D’agostino Pearson test and results were presented as volcano plots with a P value<0.01 adjusted by Holm-Sidak method and corrected for unequal variances using Welch’s correction. For frequency analysis, data were organized in contingency tables and differences were tested by chi-square test. Results were presented as mean ± standard error of the mean (SEM) for all experiments. After adjusting for multiple comparisons, p values<0.05 were considered statistically significant.

### Data, code, and materials availability

Upon publication in a peer-reviewed journal, sequencing data will be made available on ENA, and code used for analysis and generating figures will be made available on GitHub. Reagents will be made available upon request from the lead author (F.T.M.).

## Acknowledgements

We thank Ann Cloos and Swetha Srinivasaraghavan for their assistance with hPSC culture and hypothalamic differentiation, Solomon Shepherd for providing images of live POMC-GFP cells, Anne White for providing anti-POMC antibodies, and Ben Jones from Imperial College London for helpful discussions. We are also grateful to Marcella Ma of the Genomics and Bioinformatics Core, which is funded by the UK Medical Research Council (MRC) Metabolic Disease Unit (MRC_MC_UU_00014/5) and a Wellcome Trust Major Award (208363/Z/17/Z), Chris Smith of the Tissue and Cell Imaging core facility for assistance with imaging, and the staff at the Genomics Core Facility of the Cancer Research UK Cambridge Institute for assistance with sequencing. We also thank the Histopathology/ISH core facility at the Institute of Metabolic Science-Metabolic Research Laboratories and Cancer Research UK-Cambridge Institute for the assistance with RNAscope.

F.T.M. is a New York Stem Cell Foundation -Robertson Investigator [NYSCF-R-156]. H-J.C.C., I.M., and A.Y. were supported by funding from the Chan Zuckerberg Initiative -Ben Barres Early Career Investigator [CZI NDCN 191942, 10.37921/429861umrcjh]. F.T.M. and S.M. and the costs of this study were supported by the Wellcome Trust and Royal Society [211221/Z/18/Z]. C.A. was supported by a Wellcome investigator award (220271/Z/20/Z), held by F.R. and F.M.G. D.S. was supported by the University of Cambridge’s Ukrainian Academic Support Scheme and a Rowan Williams Cambridge Studentship from the Cambridge Trust. We are grateful for the support of the David James Trust. For open access, the authors have applied a CC-BY public copyright license to any Author Accepted Manuscript version arising from this submission.

## Author contributions

The study was conceptualized by S.M. and F.T.M., who wrote the manuscript with the help of H.-J.C.C. and A.Y., and reviewed and edited contributions from all authors. F.T.M. acquired funding to support this work, administered the project, and supervised studies together with J.M. Data were collected and analyzed by S.M., H.-J.C.C., O.C., A.Y., C.A.A., and F.T.M., and visualized by S.M., H.-J.C.C., A.Y., and F.T.M. D.S., J.C., V.M., and I.M. helped generate cells and reagents, and provided intellectual contributions. J.M., F.G., J.A.T. and F.R. provided technical advice and intellectual contributions. Underlying methodologies were developed by S.M., H.-J.C.C., and O.C.

## Inclusion and diversity

We support inclusive, diverse, and equitable conduct of research.

## Supplementary tables

**Table S1. Quantification of GLP1R puncta and gene expression in POMC reporter cell lines. A)** Quantification of GLP1R puncta per neuron was performed in both POMC-GFP and POMC-NeonGreen reporter cell lines. Cells with POMC mRNA showed significantly more GLP1R puncta compared with POMC-neurons in both lines (Mann–Whitney U test, p<0.05). Data are presented as mean ± 95% CI and median with Interquartile Range (IQR) (POMC-/GLP1R+ n=5,870, POMC+/GLP1R+ n=353 cells for POMC-GFP cell line; POMC-/GLP1R+ n=496, POMC+/GLP1R+ n=93 for POMC-NeonGreen cell line). **B,C)** Counts of puncta for single-neuron for POMC-GFP (B) and POMC-NG (C). Each row represents an individual segmented neuron (Object ID) along with the number of detected puncta (Copies), fluorescent area (µm²) and total cell area (µm²). Zeros indicate absence of detectable fluorescence above threshold in that channel. FITC and Cy3 correspond to fluorescence signals used to identify POMC and GLP1R expression, respectively. **D)** Bulk RNA sequencing of FACS-purified cells derived from a hypothalamic differentiation of a POMC-GFP reporter cell line into GFP-(n=2) and GFP+ (n=6) cell populations. Each row displays the gene ID and gene expression levels in log counts per million (LogCPM). **E)** Number and percentage of cells in each of 10 clusters of hPSC-derived hypothalamic cells showing cells that express *GLP1R* (≥1 UMI).

**Table S2. Calcium imaging responses of human hypothalamic cells to GLP-1R agonists. A)** <u>Top table</u>: Summary of GFP- or POMC-GFP+ neuron responses to GLP-1, semaglutide, liraglutide, tirzepatide, exendin-D3 and exendin-F1, as well as POMC-NG+ or NG-neuron responses to semaglutide. AUC_response_ and AUC_baseline_ were calculated respectively during three-minute time windows before and after two minutes of 200 nM agonist administration or agonists/antagonist co-administration. Statistical significance between median AUC responses recorded in GFP- or in POMC-GFP+ neurons was calculated for each agonist by unpaired, non-parametric Mann-Whitney t-test. <u>Second table</u>: POMC-GFP+ neurons show significantly higher AUC responses (AUC_response_ - AUC_baseline_) than GFP-neurons to the GLP-1R agonists GLP-1 (p<0.0001), semaglutide (p<0.0001) and liraglutide (p<0.05) but not to the biased agonists exendin-D3, exendin-F1 and the GIPR/GLP-1R dual agonist tirzepatide (p>0.05). Co-administration of equimolar ratios of GLP-1. <u>Third table</u>: and the GLP-1R antagonist exendin-(9-39) significantly reduced (p<0.0001) neuronal responses for GLP-1, and an excess of exendin-(9-39) (1 𝝁M) was sufficient to completely block (p<0.0001) POMC-GFP+ neuron responses to 30 nM semaglutide. <u>Fourth table</u>: Number of responding and non-responding neurons within POMC-GFP+ and GFP− or for POMC-NG+ and NG− populations, along with total cell numbers analyzed per condition. Neurons were classified as responders or non-responders based on calcium imaging response criteria defined in the Methods. Differences in the proportion of responding neurons between POMC-GFP+ and GFP− populations were assessed using chi-square tests, with *p*<0.05 considered statistically significant. Statistical significance is reported for each agonist. **B-F)** Summary data for POMG-GFP+ and GFP-neurons that significantly responded to GLP-1 (B), and underlying normalized calcium indicator dye fluorescence intensity values (ΔF/F_0_) by acquisition time point (in minutes) for recordings from individual neurons organized by POMG-GFP+ responders (C) and non-responders (D), and GFP-responders (E) and non-responders (F). Time windows used for the calculation of the AUC are highlighted in yellow for the baseline and in orange for the responses. **G-AO)** Tables organized as in B-F but for semaglutide (G-K), liraglutide (L-P), tirzepatide (Q-U), GLP-1 with and without exendin-(9-39) (V-X), semaglutide with and without exendin-(9-39) (Y,Z), exendin-D3 (AA-AE), exendin-F1 (AF-AJ) and for semaglutide treatment of the POMC-NG cell line (AK-AO).

**Table S3. Electrophysiological responses of POMC neurons differentiated from POMC-GFP and POMC-NG IPSCs lines to semaglutide. A)** Average recorded membrane potential (POMC-GFP+, n=9; POMC-NG+, n=8) and action potential firing rate (POMC-diGFP+, n=7; POMC-NG+, n=7) per minute during three-minute time windows before and after administration of 200 nM semaglutide. Data are displayed as Mean (mV) with 95% CI and Median (mV) with interquartile ranges (IQR, Q1-Q3). Statistical significance (p<0.05) between median recorded membrane potential and action potential firing rate per minute before and after the application of semaglutide were calculated by paired, Non-Parametric Wilcoxon matched-pairs signed rank test. **B)** Instantaneous frequency (Hz) recorded from n=4 POMC+ neurons under baseline conditions and after exposure to 200 nM semaglutide. The data include the number of measurements, median values with interquartile ranges (IQR, Q1-Q3), and the median of differences with 95% CI. Statistical significance between baseline and post-sema conditions was calculated using an unpaired, non-parametric Mann–Whitney test. **C)** Significantly up- or downregulated genes (|log2FC|≥0.5) in purified GFP+ neurons between vehicle- and semaglutide-treated groups (n=6 technical replicates per group) are highlighted in light orange. A more stringent subset of these genes that reached statistical significance at FDR<0.05 were classified as differentially expressed genes (dark orange) between vehicle- and semaglutide-treated neurons. **D,E**) Results of KEGG gene ontology enrichment analysis on genes significantly downregulated (D) or upregulated (E) in response to semaglutide.

**Table S4. Effect of VGCC blockers on the semaglutide responses recorded in human hypothalamic cells. A)** Effects of semaglutide and nifedipine on membrane potential and firing activity in POMC neurons. Electrophysiological recordings from POMC-GFP+ and POMC-NG+ neurons showing changes in membrane potential and firing frequency in response to 200 nM semaglutide and 20 μM nifedipine. Semaglutide depolarized membrane potential and increased action potential frequency in neurons relative to baseline. Successive application of the L-type VGCC blocker nifedipine attenuated both depolarization and firing activity. Data represent mean 95% CI, median [IQR], and median of differences. Statistical significance between semaglutide and nifedipine conditions was assessed using a paired, non-parametric Wilcoxon matched-pairs signed-rank test. **B)** Summary of calcium imaging data showing the effects of VGCC blockers on neurons responding to semaglutide. Averaged calcium responses were calculated as AUCs at baseline and after perfusion with semaglutide for 120 seconds, after which responses were allowed to plateau before the addition of a VGCCs blocker for 8 minutes. Mean and median AUC values (95% CI, IQR) are presented for semaglutide responses before and after application of VGCCs blockers. Statistical analysis was performed using a paired, non-parametric Wilcoxon matched-pairs signed-rank test. **C-G)** Single neuron AUCs calculated during two-minute time windows at baseline, after semaglutide application (pre-VGCC blocker), and following VGCC inhibition (post-VGCC blocker) with ω-Agatoxin (C), TTA-P2 (D), benidipine (E), nifedipine (F) and co-application of ω-Agatoxin + TTA-P2 + benidipine (G). **H-L)** Time course and respective ΔF/F_0_ values for POMC-GFP+ neurons (highlighted in green) and GFP-neurons stimulated initially with semaglutide and then with the VGCC blockers ω-Agatoxin (H), TTA-P2 (I), benidipine (J), nifedipine (K) and combined ω-Agatoxin + TTA-P2 + benidipine (L). Time windows used for the calculation of the AUCs are highlighted in yellow for the baseline and in orange for the response to semaglutide, and in blue for the effect of the blocker.

## References

1. Acuna-Goycolea, Claudio, and Anthony van den Pol. 2004. “Glucagon-like Peptide 1 Excites Hypocretin/orexin Neurons by Direct and Indirect Mechanisms: Implications for Viscera-Mediated Arousal.” The Journal of Neuroscience: The Official Journal of the Society for Neuroscience 24 (37): 8141–8152.

2. Alhadeff, Amber L., Laura E. Rupprecht, and Matthew R. Hayes. 2012. “GLP-1 Neurons in the Nucleus of the Solitary Tract Project Directly to the Ventral Tegmental Area and Nucleus Accumbens to Control for Food Intake.” Endocrinology 153 (2): 647–658.

3. Baldini, Giulia, and Kevin D. Phelan. 2019. “The Melanocortin Pathway and Control of Appetite-Progress and Therapeutic Implications.” The Journal of Endocrinology 241 (1): R1–R33.

4. Bessesen, Daniel H., and Luc F. Van Gaal. 2018. “Progress and Challenges in Anti-Obesity Pharmacotherapy.” The Lancet. Diabetes & Endocrinology 6 (3): 237–248.

5. Brennecke, Philip, Simon Anders, Jong Kyoung Kim, et al. 2013. “Accounting for Technical Noise in Single-Cell RNA-Seq Experiments.” Nature Methods 10 (11): 1093–1095.

6. Budde, Thomas, Sven Meuth, and Hans-Christian Pape. 2002. “Calcium-Dependent Inactivation of Neuronal Calcium Channels.” Nature Reviews Neuroscience 3 (11): 873–883.

7. Catterall, William A. 2011. “Voltage-Gated Calcium Channels.” Cold Spring Harbor Perspectives in Biology 3 (8): a003947.

8. Chen, Hsiao-Jou Cortina, Simone Mazzaferro, Tian Tian, Iman Mali, and Florian T. Merkle. 2023. “Differentiation, Transcriptomic Profiling, and Calcium Imaging of Human Hypothalamic Neurons.” Current Protocols 3 (6): e786.

9. Chen, Hsiao-Jou Cortina, Andrian Yang, Simone Mazzaferro, et al. 2023. “Profiling Human Hypothalamic Neurons Reveals a Candidate Combination Drug Therapy for Weight Loss.” In bioRxiv. July 19. 10.1101/2023.07.18.549357.

10. Coe, H., and M. Michalak. 2009. “Calcium Binding Chaperones of the Endoplasmic Reticulum.” General Physiology and Biophysics 28 Spec No Focus. https://pubmed.ncbi.nlm.nih.gov/20093733/.

11. Cone, Roger D. 2005. “Anatomy and Regulation of the Central Melanocortin System.” Nature Neuroscience 8 (5): 571–578.

12. Coskun, Tamer, Kyle W. Sloop, Corina Loghin, et al. 2018. “LY3298176, a Novel Dual GIP and GLP-1 Receptor Agonist for the Treatment of Type 2 Diabetes Mellitus: From Discovery to Clinical Proof of Concept.” Molecular Metabolism 18 (December): 3–14.

13. Coskun, Tamer, Shweta Urva, William C. Roell, et al. 2022. “LY3437943, a Novel Triple Glucagon, GIP, and GLP-1 Receptor Agonist for Glycemic Control and Weight Loss: From Discovery to Clinical Proof of Concept.” Cell Metabolism 34 (9): 1234–1247.e9.

14. Cowley, M. A., J. L. Smart, M. Rubinstein, et al. 2001. “Leptin Activates Anorexigenic POMC Neurons through a Neural Network in the Arcuate Nucleus.” Nature 411 (6836): 480–484.

15. Dittmann, Marie T., Gabriella Lakatos, Jodie F. Wainwright, et al. 2024. “Low Resting Metabolic Rate and Increased Hunger due to β-MSH and β-Endorphin Deletion in a Canine Model.” Science Advances 10 (10): eadj3823.

16. Dobin, A., C. A. Davis, F. Schlesinger, et al. 2013. “STAR: Ultrafast Universal RNA-Seq Aligner.” Bioinformatics 29 (1). 10.1093/bioinformatics/bts635.

17. Dong, Yanbin, Jamie Carty, Nitsan Goldstein, et al. 2021. “Time and Metabolic State-Dependent Effects of GLP-1R Agonists on NPY/AgRP and POMC Neuronal Activity in Vivo.” Molecular Metabolism 54 (December): 101352.

18. During, Matthew J., Lei Cao, David S. Zuzga, et al. 2003. “Glucagon-like Peptide-1 Receptor Is Involved in Learning and Neuroprotection.” Nature Medicine 9 (9): 1173–1179.

19. Fang, Zijian, Shiqian Chen, Philip Pickford, et al. 2020. “The Influence of Peptide Context on Signaling and Trafficking of Glucagon-like Peptide-1 Receptor Biased Agonists.” ACS Pharmacology & Translational Science 3 (2): 345.

20. Farooqi, I. Sadaf, Julia M. Keogh, Giles S. H. Yeo, Emma J. Lank, Tim Cheetham, and Stephen O’Rahilly. 2003. “Clinical Spectrum of Obesity and Mutations in the Melanocortin 4 Receptor Gene.” The New England Journal of Medicine 348 (12): 1085–1095.

21. Farooqi, Sadaf, and Stephen O’Rahilly. 2006. “Genetics of Obesity in Humans.” Endocrine Reviews 27 (7): 710–718.

22. Fenselau, Henning, John N. Campbell, Anne M. J. Verstegen, et al. 2017. “A Rapidly Acting Glutamatergic ARC→PVH Satiety Circuit Postsynaptically Regulated by α-MSH.” Nature Neuroscience 20 (1): 42–51.

23. Ford, Neil C., and Mark L. Baccei. 2016. “Inward-Rectifying K+ (Kir2) Leak Conductance Dampens the Excitability of Lamina I Projection Neurons in the Neonatal Rat.” Neuroscience 339 (October): 502.

24. Gabery, Sanaz, Casper G. Salinas, Sarah J. Paulsen, et al. 2020. “Semaglutide Lowers Body Weight in Rodents via Distributed Neural Pathways.” JCI Insight 5 (6). 10.1172/jci.insight.133429.

25. Goldspink, Deborah A., Van B. Lu, Emily L. Miedzybrodzka, et al. 2020. “Labeling and Characterization of Human GLP-1-Secreting L-Cells in Primary Ileal Organoid Culture.” Cell Reports 31 (13): 107833.

26. Gomez, Edith, Catrin Pritchard, and Terence P. Herbert. 2002. “cAMP-Dependent Protein Kinase and Ca2+ Influx through L-Type Voltage-Gated Calcium Channels Mediate Raf-Independent Activation of Extracellular Regulated Kinase in Response to Glucagon-like Peptide-1 in Pancreatic Beta-Cells.” The Journal of Biological Chemistry 277 (50): 48146–48151.

27. Hael, C. E., D. Rojo, D. P. Orquera, M. J. Low, and M. Rubinstein. 2020. “The Transcriptional Regulator PRDM12 Is Critical for Pomc Expression in the Mouse Hypothalamus and Controlling Food Intake, Adiposity, and Body Weight.” Molecular Metabolism 34 (April). 10.1016/j.molmet.2020.01.007.

28. Halawi, Houssam, Disha Khemani, Deborah Eckert, et al. 2017. “Effects of Liraglutide on Weight, Satiation, and Gastric Functions in Obesity: A Randomised, Placebo-Controlled Pilot Trial.” *The Lancet*. Gastroenterology & Hepatology 2 (12): 890–899.

29. Hayes, Matthew R., Theresa M. Leichner, Shiru Zhao, et al. 2011. “Intracellular Signals Mediating the Food Intake-Suppressive Effects of Hindbrain Glucagon-like Peptide-1 Receptor Activation.” Cell Metabolism 13 (3): 320–330.

30. He, Zhenyan, Yong Gao, Linh Lieu, et al. 2019. “Direct and Indirect Effects of Liraglutide on Hypothalamic POMC and NPY/AgRP Neurons - Implications for Energy Balance and Glucose Control.” Molecular Metabolism 28 (October): 120–134.

31. Hibino, Hiroshi, Atsushi Inanobe, Kazuharu Furutani, Shingo Murakami, Ian Findlay, and Yoshihisa Kurachi. 2010. “Inwardly Rectifying Potassium Channels: Their Structure, Function, and Physiological Roles.” *Physiological Reviews*, ahead of print, January 1. 10.1152/physrev.00021.2009.

32. Huang, Kuei-Pin, Alisha A. Acosta, Misgana Y. Ghidewon, et al. 2024. “Dissociable Hindbrain GLP1R Circuits for Satiety and Aversion.” Nature 632 (8025): 585–593.

33. Huguenard, J. R. 1996. “Low-Threshold Calcium Currents in Central Nervous System Neurons.” Annual Review of Physiology 58 (1): 329–348.

34. Huszar, D., C. A. Lynch, V. Fairchild-Huntress, et al. 1997. “Targeted Disruption of the Melanocortin-4 Receptor Results in Obesity in Mice.” Cell 88 (1): 131–141.

35. Irannejad, Roshanak, Jin C. Tomshine, Jon R. Tomshine, et al. 2013. “Conformational Biosensors Reveal GPCR Signalling from Endosomes.” Nature 495 (7442): 534–538.

36. Jastreboff, Ania M., Louis J. Aronne, Nadia N. Ahmad, et al. 2022. “Tirzepatide Once Weekly for the Treatment of Obesity.” The New England Journal of Medicine 387 (3): 205–216.

37. Jones, B. 2022. “The Therapeutic Potential of GLP-1 Receptor Biased Agonism.” British Journal of Pharmacology 179 (4). 10.1111/bph.15497.

38. Jones, Ben, Teresa Buenaventura, Nisha Kanda, et al. 2018. “Targeting GLP-1 Receptor Trafficking to Improve Agonist Efficacy.” Nature Communications 9 (April): 1602.

39. Kanehisa, Minoru, Miho Furumichi, Yoko Sato, Masayuki Kawashima, and Mari Ishiguro-Watanabe. 2023. “KEGG for Taxonomy-Based Analysis of Pathways and Genomes.” Nucleic Acids Research 51 (D1): D587–D592.

40. Kawatani, Masahiro, Yuichiro Yamada, and Masahito Kawatani. 2018. “Glucagon-like Peptide-1 (GLP-1) Action in the Mouse Area Postrema Neurons.” Peptides 107 (September): 68–74.

41. Kemp, Paul J., David J. Rushton, Polina L. Yarova, et al. 2016. “Improving and Accelerating the Differentiation and Functional Maturation of Human Stem Cell-Derived Neurons: Role of Extracellular Calcium and GABA.” The Journal of Physiology 594 (22): 6583–6594.

42. Kirwan, Peter, Magdalena Jura, and Florian T. Merkle. 2017. “Generation and Characterization of Functional Human Hypothalamic Neurons.” Current Protocols in Neuroscience / Editorial Board, Jacqueline N. Crawley … [et Al.] 81 (October): 3.33.1–3.33.24.

43. Kirwan, Peter, Richard G. Kay, Bas Brouwers, et al. 2018. “Quantitative Mass Spectrometry for Human Melanocortin Peptides in Vitro and in Vivo Suggests Prominent Roles for β-MSH and Desacetyl α-MSH in Energy Homeostasis.” Molecular Metabolism 17 (November): 82–97.

44. Klaauw, Agatha A. van der, Sophie Croizier, Edson Mendes de Oliveira, et al. 2019. “Human Semaphorin 3 Variants Link Melanocortin Circuit Development and Energy Balance.” Cell 176 (4): 729–742.e18.

45. Knerr, Patrick J., Stephanie A. Mowery, Jonathan D. Douros, et al. 2022. “Next Generation GLP-1/GIP/glucagon Triple Agonists Normalize Body Weight in Obese Mice.” Molecular Metabolism 63 (September): 101533.

46. Knudsen, Lotte Bjerre, and Jesper Lau. 2019. “The Discovery and Development of Liraglutide and Semaglutide.” Frontiers in Endocrinology 10 (April): 155.

47. Kopp, Katherine O., Elliot J. Glotfelty, Yazhou Li, and Nigel H. Greig. 2022. “Glucagon-like Peptide-1 (GLP-1) Receptor Agonists and Neuroinflammation: Implications for Neurodegenerative Disease Treatment.” Pharmacological Research: The Official Journal of the Italian Pharmacological Society 186 (December): 106550.

48. Krude, H., H. Biebermann, W. Luck, R. Horn, G. Brabant, and A. Grüters. 1998. “Severe Early-Onset Obesity, Adrenal Insufficiency and Red Hair Pigmentation Caused by POMC Mutations in Humans.” Nature Genetics 19 (2): 155–157.

49. Lau, Jesper, Paw Bloch, Lauge Schäffer, et al. 2015. “Discovery of the Once-Weekly Glucagon-Like Peptide-1 (GLP-1) Analogue Semaglutide.” Journal of Medicinal Chemistry 58 (18): 7370–7380.

50. Lee, Yung Seng, Ben G. Challis, Darren A. Thompson, et al. 2006. “A POMC Variant Implicates Beta-Melanocyte-Stimulating Hormone in the Control of Human Energy Balance.” Cell Metabolism 3 (2): 135–140.

51. Liao, Yang, Gordon K. Smyth, and Wei Shi. 2014. “featureCounts: An Efficient General Purpose Program for Assigning Sequence Reads to Genomic Features.” Bioinformatics 30 (7): 923–930.

52. Lippiat, Jonathan D. 2008. “Whole-Cell Recording Using the Perforated Patch Clamp Technique.” Methods in Molecular Biology 491: 141–149.

53. Littleton, Sheridan H., Khanh B. Trang, Christina M. Volpe, et al. 2024. “Variant-to-Function Analysis of the Childhood Obesity chr12q13 Locus Implicates rs7132908 as a Causal Variant within the 3′ UTR of FAIM2.” Cell Genomics 4 (5): 100556.

54. Li, Yazhou, Tracyann Perry, Mark S. Kindy, et al. 2009. “GLP-1 Receptor Stimulation Preserves Primary Cortical and Dopaminergic Neurons in Cellular and Rodent Models of Stroke and Parkinsonism.” Proceedings of the National Academy of Sciences of the United States of America 106 (4): 1285–1290.

55. Locke, Adam E., Bratati Kahali, Sonja I. Berndt, et al. 2015. “Genetic Studies of Body Mass Index Yield New Insights for Obesity Biology.” Nature 518 (7538): 197–206.

56. Loos, Ruth J. F., and Giles S. H. Yeo. 2022. “The Genetics of Obesity: From Discovery to Biology.” Nature Reviews. Genetics 23 (2): 120–133.

57. Lucey, M., P. Pickford, S. Bitsi, et al. 2020. “Disconnect between Signalling Potency and in Vivo Efficacy of Pharmacokinetically Optimised Biased Glucagon-like Peptide-1 Receptor Agonists.” Molecular Metabolism 37 (July). 10.1016/j.molmet.2020.100991.

58. Meier, Juris J. 2012. “GLP-1 Receptor Agonists for Individualized Treatment of Type 2 Diabetes Mellitus.” Nature Reviews. Endocrinology 8 (12): 728–742.

59. Merkle, Florian T., Asif Maroof, Takafumi Wataya, et al. 2015. “Generation of Neuropeptidergic Hypothalamic Neurons from Human Pluripotent Stem Cells.” Development 142 (4): 633–643.

60. Müller, T. D., B. Finan, S. R. Bloom, et al. 2019. “Glucagon-like Peptide 1 (GLP-1).” Molecular Metabolism 30 (December): 72–130.

61. Nakajima, Kazuo, Mizuho Ishiwata, Adam Z. Weitemier, et al. 2021. “Brain-Specific Heterozygous Loss-of-Function of ATP2A2, Endoplasmic Reticulum Ca2+ Pump Responsible for Darier’s Disease, Causes Behavioral Abnormalities and a Hyper-Dopaminergic State.” Human Molecular Genetics 30 (18): 1762.

62. NCD Risk Factor Collaboration (NCD-RisC). 2024. “Worldwide Trends in Underweight and Obesity from 1990 to 2022: A Pooled Analysis of 3663 Population-Representative Studies with 222 Million Children, Adolescents, and Adults.” The Lancet 403 (10431): 1027–1050.

63. Nelson, A. B., A. H. Gittis, and S. du Lac. 2005. “Decreases in CaMKII Activity Trigger Persistent Potentiation of Intrinsic Excitability in Spontaneously Firing Vestibular Nucleus Neurons.” Neuron 46 (4). 10.1016/j.neuron.2005.04.009.

64. Okonechnikov, Konstantin, Ana Conesa, and Fernando García-Alcalde. 2016. “Qualimap 2: Advanced Multi-Sample Quality Control for High-Throughput Sequencing Data.” Bioinformatics 32 (2): 292–294.

65. Orquera, Daniela P., M. Belén Tavella, Flavio S. J. de Souza, Sofía Nasif, Malcolm J. Low, and Marcelo Rubinstein. 2019. “The Homeodomain Transcription Factor NKX2.1 Is Essential for the Early Specification of Melanocortin Neuron Identity and Activates Pomc Expression in the Developing Hypothalamus.” The Journal of Neuroscience 39 (21): 4023.

66. Panagaki, Theodora, Simon Gengler, and Christian Hölscher. 2018. “The Novel DA-CH3 Dual Incretin Restores Endoplasmic Reticulum Stress and Autophagy Impairments to Attenuate Alzheimer-Like Pathology and Cognitive Decrements in the APPSWE/PS1ΔE9 Mouse Model.” Journal of Alzheimer’s Disease: JAD 66 (1): 195–218.

67. Pantazis, C. B., A. Yang, E. Lara, et al. 2022. “A Reference Human Induced Pluripotent Stem Cell Line for Large-Scale Collaborative Studies.” Cell Stem Cell 29 (12). 10.1016/j.stem.2022.11.004.

68. Perez-Reyes, E. 2003. “Molecular Physiology of Low-Voltage-Activated T-Type Calcium Channels.” Physiological Reviews 83 (1). 10.1152/physrev.00018.2002.

69. Péterfi, Zoltán, Anett Szilvásy-Szabó, Erzsébet Farkas, et al. 2021. “Glucagon-Like Peptide-1 Regulates the Proopiomelanocortin Neurons of the Arcuate Nucleus Both Directly and Indirectly via Presynaptic Action.” Neuroendocrinology 111 (10): 986–997.

70. Qiu, J., E. J. Wagner, O. K. Rønnekleiv, and M. J. Kelly. 2018. “Insulin and Leptin Excite Anorexigenic pro-Opiomelanocortin Neurones via Activation of TRPC5 Channels.” Journal of Neuroendocrinology 30 (2). 10.1111/jne.12501.

71. Quarta, Carmelo, Marc Claret, Lori M. Zeltser, et al. 2021. “POMC Neuronal Heterogeneity in Energy Balance and beyond: An Integrated View.” Nature Metabolism 3 (3): 299–308.

72. Raudvere, Uku, Liis Kolberg, Ivan Kuzmin, et al. 2019. “g:Profiler: A Web Server for Functional Enrichment Analysis and Conversions of Gene Lists (2019 Update).” Nucleic Acids Research 47 (W1): W191–W198.

73. Reich, Niklas, and Christian Hölscher. 2022. “The Neuroprotective Effects of Glucagon-like Peptide 1 in Alzheimer’s and Parkinson’s Disease: An in-Depth Review.” Frontiers in Neuroscience 16 (September): 970925.

74. Rønnekleiv, Oline K., Yuan Fang, Chunguang Zhang, Casey C. Nestor, Peizhong Mao, and Martin J. Kelly. 2014. “Research Resource: Gene Profiling of G Protein-Coupled Receptors in the Arcuate Nucleus of the Female.” Molecular Endocrinology 28 (8): 1362–1380.

75. Secher, Anna, Jacob Jelsing, Arian F. Baquero, et al. 2014. “The Arcuate Nucleus Mediates GLP-1 Receptor Agonist Liraglutide-Dependent Weight Loss.” The Journal of Clinical Investigation 124 (10): 4473–4488.

76. Shilleh, Ali H., Katrina Viloria, Johannes Broichhagen, Jonathan E. Campbell, and David J. Hodson. 2024. “GLP1R and GIPR Expression and Signaling in Pancreatic Alpha Cells, Beta Cells and Delta Cells.” Peptides 175 (February): 171179.

77. Singh, Ishnoor, Le Wang, Baijuan Xia, et al. 2022. “Activation of Arcuate Nucleus Glucagon-like Peptide-1 Receptor-Expressing Neurons Suppresses Food Intake.” Cell & Bioscience 12 (1): 178.

78. Smith, Mark A., Kazunari Hisadome, Hind Al-Qassab, Helen Heffron, Dominic J. Withers, and Michael L. J. Ashford. 2006. “Melanocortins and Agouti-Related Protein Modulate the Excitability of Two Arcuate Nucleus Neuron Populations by Alteration of Resting Potassium Conductances.” The Journal of Physiology 578 (Pt 2): 425.

79. Sonoda, Noriyuki, Takeshi Imamura, Takeshi Yoshizaki, Jennie L. Babendure, Juu-Chin Lu, and Jerrold M. Olefsky. 2008. “Beta-Arrestin-1 Mediates Glucagon-like Peptide-1 Signaling to Insulin Secretion in Cultured Pancreatic Beta Cells.” Proceedings of the National Academy of Sciences of the United States of America 105 (18): 6614–6619.

80. Steuernagel, Lukas, Brian Y. H. Lam, Paul Klemm, et al. 2022. “HypoMap-a Unified Single-Cell Gene Expression Atlas of the Murine Hypothalamus.” Nature Metabolism 4 (10): 1402–1419.

81. Tadross, John A., Lukas Steuernagel, Georgina K. C. Dowsett, et al. 2025. “A Comprehensive Spatio-Cellular Map of the Human Hypothalamus.” Nature 639 (8055): 708–716.

82. Tan, Qiming, Seun E. Akindehin, Camila E. Orsso, et al. 2022. “Recent Advances in Incretin-Based Pharmacotherapies for the Treatment of Obesity and Diabetes.” Frontiers in Endocrinology 13 (March): 838410.

83. Vadini, Francesco, Paola G. Simeone, Andrea Boccatonda, et al. 2020. “Liraglutide Improves Memory in Obese Patients with Prediabetes or Early Type 2 Diabetes: A Randomized, Controlled Study.” International Journal of Obesity 44 (6): 1254–1263.

84. Wilding, John P. H., Rachel L. Batterham, Salvatore Calanna, et al. 2021. “Once-Weekly Semaglutide in Adults with Overweight or Obesity.” The New England Journal of Medicine 384 (11): 989–1002.

85. Wu, Peipei, Yuxiao Zhao, Xianghua Zhuang, Aili Sun, Yuan Zhang, and Yihong Ni. 2018. “Low Glucagon-like Peptide-1 (GLP-1) Concentration in Serum Is Indicative of Mild Cognitive Impairment in Type 2 Diabetes Patients.” Clinical Neurology and Neurosurgery 174 (November): 203–206.

86. Yaswen, L., N. Diehl, M. B. Brennan, and U. Hochgeschwender. 1999. “Obesity in the Mouse Model of pro-Opiomelanocortin Deficiency Responds to Peripheral Melanocortin.” Nature Medicine 5 (9): 1066–1070.

87. Yeo, Giles S. H., Chiao-Chien Connie Hung, Justin Rochford, et al. 2004. “A de Novo Mutation Affecting Human TrkB Associated with Severe Obesity and Developmental Delay.” Nature Neuroscience 7 (11): 1187–1189.

88. Yeo, Giles S. H., Emma J. Lank, I. Sadaf Farooqi, Julia Keogh, Benjamin G. Challis, and Stephen O’Rahilly. 2003. “Mutations in the Human Melanocortin-4 Receptor Gene Associated with Severe Familial Obesity Disrupts Receptor Function through Multiple Molecular Mechanisms.” Human Molecular Genetics 12 (5): 561–574.

89. Yu, Hui, Marcelo Rubinstein, and Malcolm J. Low. 2022. Developmental Single-Cell Transcriptomics of Hypothalamic POMC Neurons Reveal the Genetic Trajectories of Multiple Neuropeptidergic Phenotypes. January 19. 10.7554/eLife.72883.

90. Zamponi, Gerald W., Joerg Striessnig, Alexandra Koschak, and Annette C. Dolphin. 2015. “The Physiology, Pathology, and Pharmacology of Voltage-Gated Calcium Channels and Their Future Therapeutic Potential.” Pharmacological Reviews 67 (4): 821–870.

91. Zhao, Fenghui, Qingtong Zhou, Zhaotong Cong, et al. 2022. “Structural Insights into Multiplexed Pharmacological Actions of Tirzepatide and Peptide 20 at the GIP, GLP-1 or Glucagon Receptors.” Nature Communications 13 (1): 1–16.

